# Huge genetic diversity of *Schizothecium tetrasporum* (Wint.). N. Lundq.: delimitation of 18 species distributed into three complexes through genome sequencing

**DOI:** 10.1101/2025.03.27.645579

**Authors:** Elsa De Filippo, Valérie Gautier, Christophe Lalanne, Emilie Levert, Elizabeth Chahine, Fanny E. Hartmann, Tatiana Giraud, Philippe Silar

## Abstract

Analyses of the genetic diversity of well-studied fungi of the *Sordariales* order, such as *Neurospora spp.* and *Podospora anserina* (syn. *Triangularia anserina*), have shown that the species classically defined by morphology are often complexes of cryptic species. Here, we report on the species delimitation among 76 strains producing mycelium and sexual reproductive structures identical to those of the pseudo-homothallic *Sordariales* species *Schizothecium tetrasporum* (syn. *Neoschizothecium tetrasporum*). Their whole genomes were sequenced as well as those of six strains closely related to *Schizothecium tetrasporum* but producing eight-spored asci instead of four-spored ones. The clustering based on the Average Nucleotide Identity (ANI) between the genomes identified eighteen species grouped into three clades, which were further supported by a phylogenetic tree constructed with whole genome Single Nucleotide Polymorphisms (SNPs). Based on their contrasting breeding systems and their large evolutionary distances, we considered the three clades as distinct species complexes. Indeed, two of them, the *Schizothecium tetrasporum* and *Schizothecium pseudotetrasporum* complexes, contains pseudo-homothallic species producing four-spored asci, while the third one, which we named *Schizothecium octosporum*, contains heterothallic species producing eight-spored asci. Surprisingly it was nestled between the two complexes of pseudo-homothallic species. Our data reveals thus a huge genetic diversity of the *Schizothecium tetrasporum* morpho-species and a convergent evolution of pseudo-homothallism or reversion to heterothallism within the complexes. An epitype for *Schizothecium tetrasporum sensus stricto* is defined and the seventeen new *Schizothecium* species are formally described.

## Introduction

DNA sequence analyses of species belonging to the *Sordariales* order (*Ascomycota*, class *Sordariomycetes*) often reveal that they actually constitute species complexes, underscoring a large genetic diversity within morphologically-homogeneous groups. This has been the case for the eight-spored *Neurospora crassa* and *Neurospora Intermedia* (Dettman et al. 2003a; Dettman et al. 2003b), *Neurospora discreta* (Dettman et al. 2006) and *Neurospora sitophila* (Svedberg et al. 2021)), as well as for the four-spored *Neurospora tetrasperma* (Menkis et al. 2009), all belonging to the *Sordariaceae* family. A total of 26 species are now recognized instead of the five initial species (Gladieux et al. 2020). Another example is *Podospora anserina*, belonging to the *Podosporaceae*, and that has been split into eight species (Boucher et al. 2017; Silar et al. 2025).

*Schizothecium tetrasporum* (Wint.) N. Lundq. is a pseudo-homothallic *Sordariales* species belonging to the *Schizotheciaceae* family. As such, it produces asci with four heterokaryotic ascospores originating from binucleated heterokaryotic progenitor cells carrying two nuclei with compatible mating types, *i.e.*, one *mat1-1* (or *mat-*) nucleus and one *mat1-2* (or *mat+*) nucleus. It also yields at low frequencies 5-, 6- or even 7-spored asci, in which some primarily binucleated heterokaryotic ascospores do not properly differentiate and are replaced with homokaryotic ones stemming from uninucleated progenitor cells. Heterokaryotic ascospores are easily differentiated from homokaryotic ones by their larger size. *S. tetrasporum* has recently been studied for its recombination suppression around the mating type locus, but has otherwise seldom been used as a model (Vittorelli et al. 2023). Therefore, there is limited knowledge on this fungus, especially on its diversity. It was originally described as *Sordaria tetraspora* by Georg Winter (Winter 1871) and later on placed in the *Schizothecium* genus by Nils Lundqvist (in (Lundqvist 1972) page 256). Lundqvist stated that no original collection exists as the first description does not mention any herbarium specimen and he also stated that a “neotype will be selected later”. Unfortunately, to the best of our knowledge, a neotype has never been selected by Lundqvist. However, a neotype was selected by Bell and Mahoney (Bell and Mahoney 1995), but it consists of a mouse dung pellet onto which “no complete perithecia or asci remain” making it poorly suitable for a type, especially in the view of the species complexes described here that can only be differentiated by extensive sequence determinations. In a controversial paper (see (Marin-Felix and Miller 2022)), Huang *et al*. renamed the fungus *Neoschizothecium tetrasporum* (G. Winter) S.K. Huang & K.D. Hyde and used the CBS394.87 strain for their phylogenetic analyses (Huang et al. 2021). They did not define any epitype for this species. Unfortunately, The CBS394.87 strain was mislabeled in the Westerdijk Institute collection and is actually a strain of *Schizothecium inaequale* (Cain) N. lundq. (correction is now made in the Westerdijk Institute collection), rendering void the phylogenetic placement of *S. tetrasporum* by Huang *et al*. (2021).

The new technologies enabling to sequence complete genomes at a low cost now permit to easily differentiate fungal species through average nucleotide identity (or ANI) calculations between assembled genome sequences (Lalanne and Silar 2024; Gostinčar 2020). Indeed, ANI calculated on whole genomes including coding and non-coding regions are usually higher than 99% (and often close to 99.9%) when comparing fungal strains from the same species, while they are lower than 99% when comparing fungal strains from different species (Lalanne and Silar 2024). Additionally, SNP calling against a reference strain can also be used to construct phylogenomic trees (Lo et al. 2023; Ropars et al. 2020; Ali et al. 2023). Here, we apply these methods based on whole genome sequences to analyze 76 four-spored strains identified morphologically as *S. tetrasporum*, including seven from culture collections and 69 strains newly collected from nature. The analyses enabled to delimit 16 distinct cryptic species. We also analyzed six eight-spored strains, including two that nested within the four-spored ones based on their ITS sequences. Genome sequences and phylogenomic trees confirmed that these two strains were indeed nested within the four-spored species and corresponded to two different species. Owing to the lack of type for *S. tetrasporum*, the controversial nature of the *Neoschizothecium* genus (Marin-Felix and Miller 2022) and the use of an incorrect strain by Huang *et al*. (2021), we propose here to redefine *S. tetrasporum* by selecting a suitable epitype for *S. tetrasporum sensu stricto*, and to describe the 17 new species identified through complete genome sequencing.

## Materials and Methods

### Strain isolation

Six strains were purchased from the Westerdijk collection (CBS strains) and one from the CABI culture collection (IMI320477 strain) (Table 1). All proved to have retained the ability to undergo sexual reproduction. However, IMI320477 only yielded unpigmented ascospores. The 68 strains with a PSN prefix and the PSQ32 strain were collected between 2018 and 2022 from dung and soil samples from various origins (Table 1). The samples were collected in Chile, Italy, Canada and New Zealand, which are not parties of the Nagoya protocol, in Denmark that does not regulate access to genetic resources within its national jurisdiction and in France, where there was an exception for micro-organisms at the time of sampling.

**Table 1.**
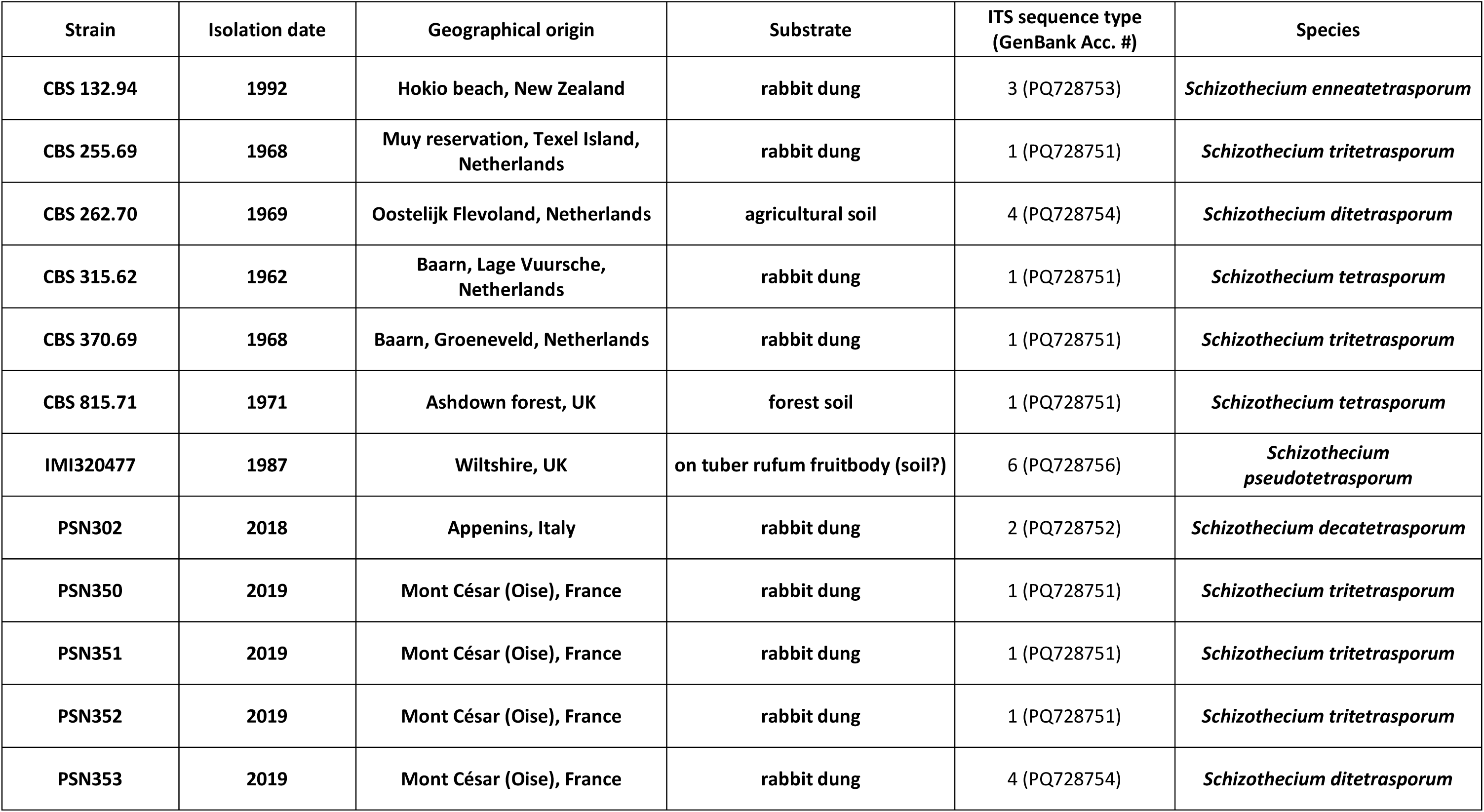
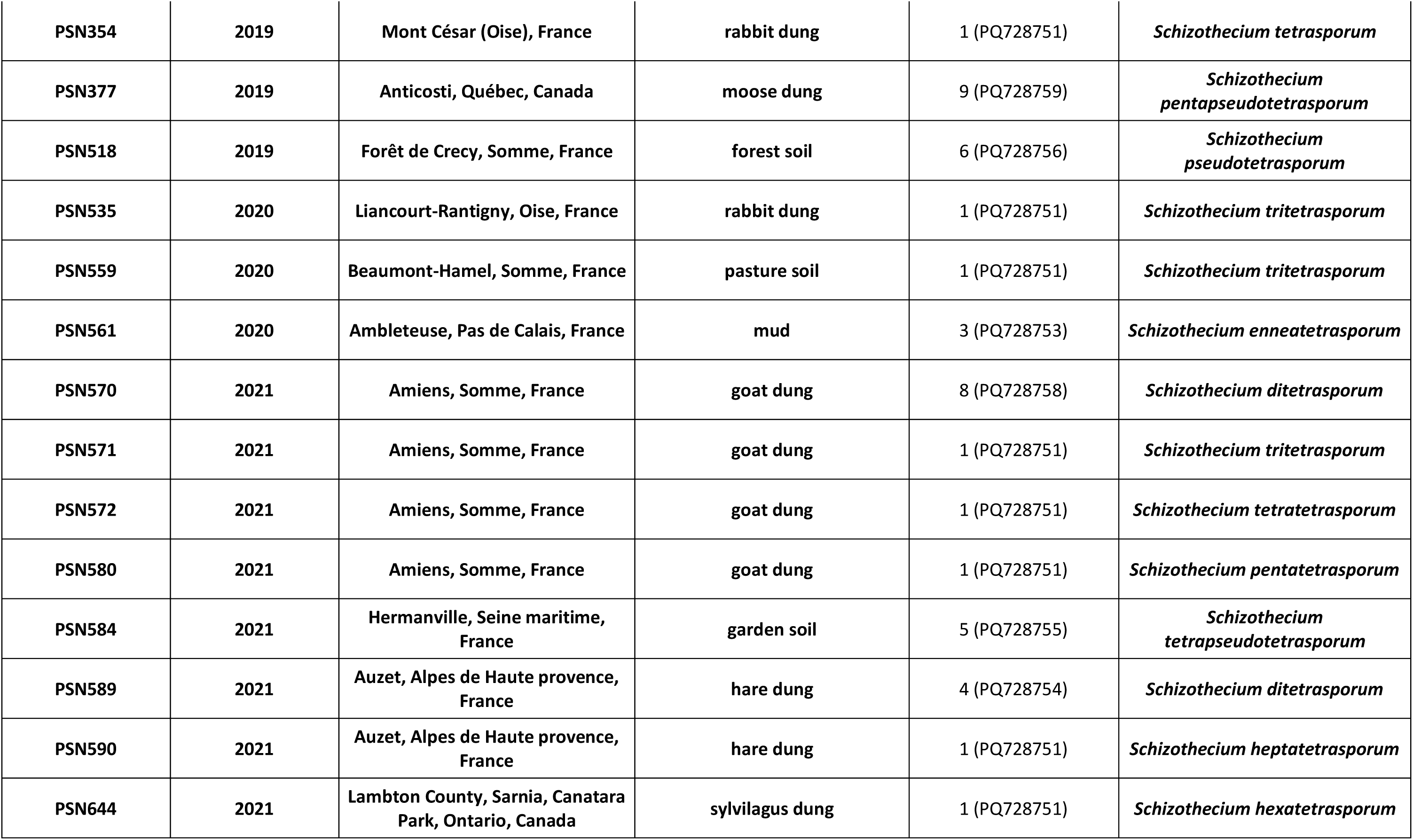
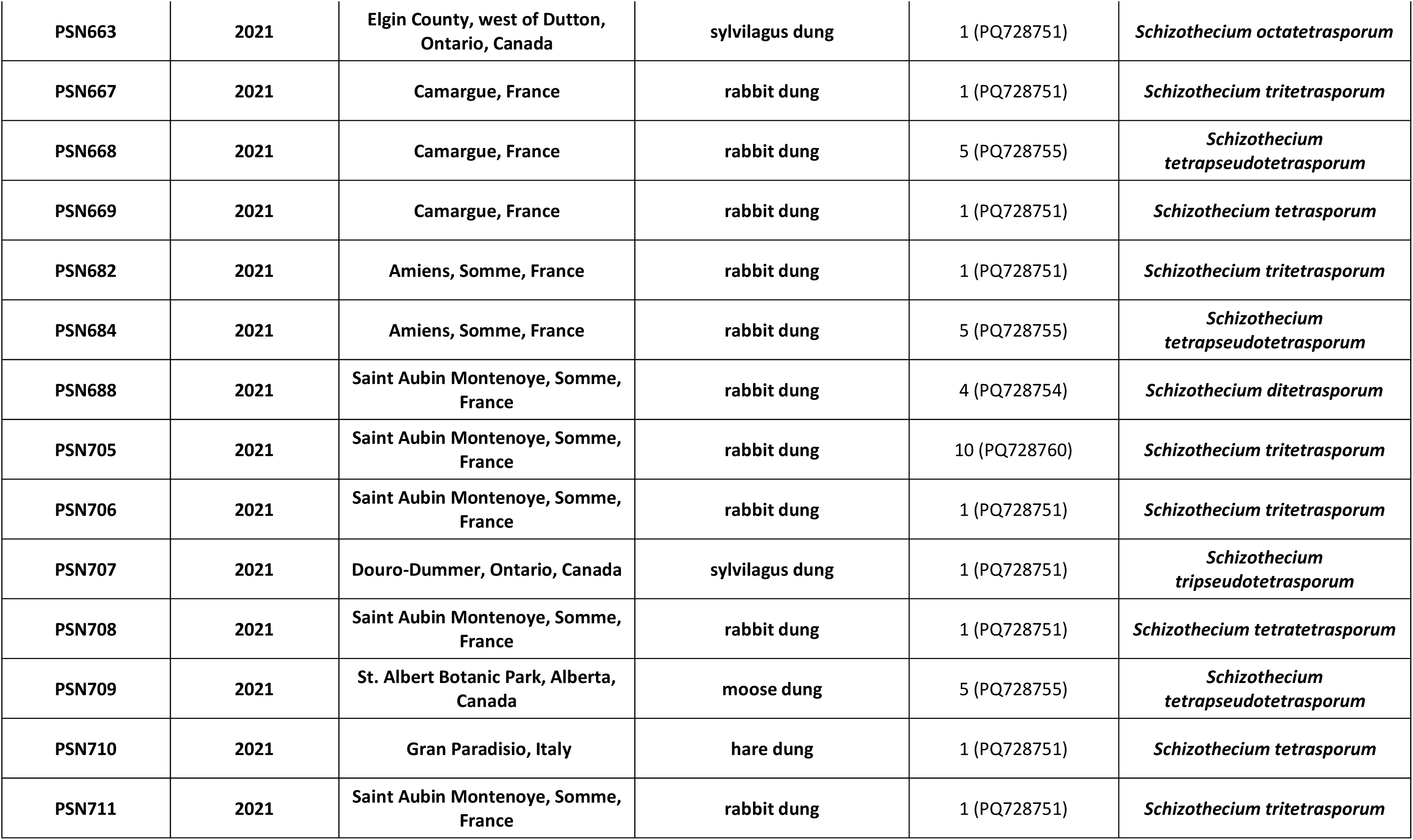
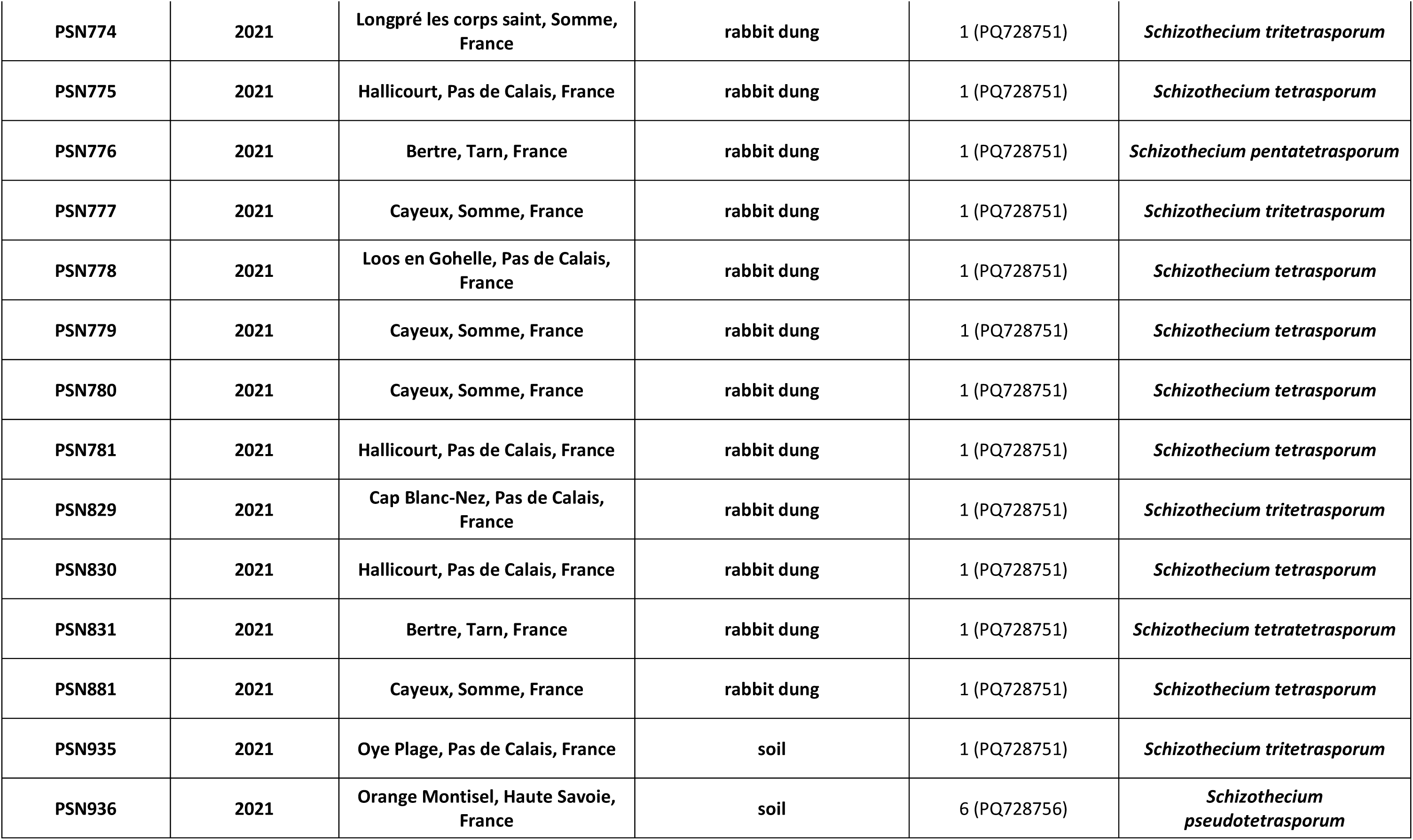
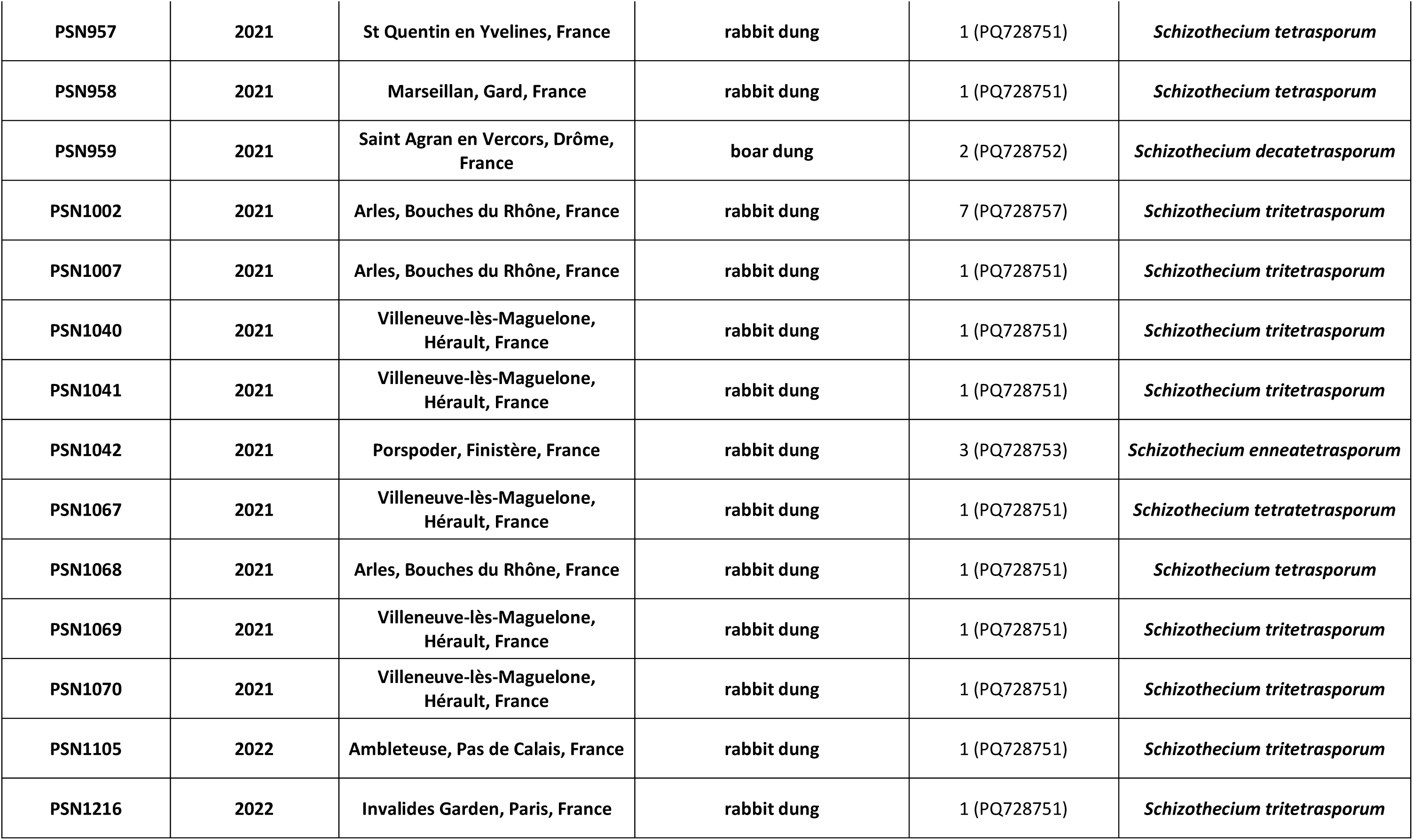
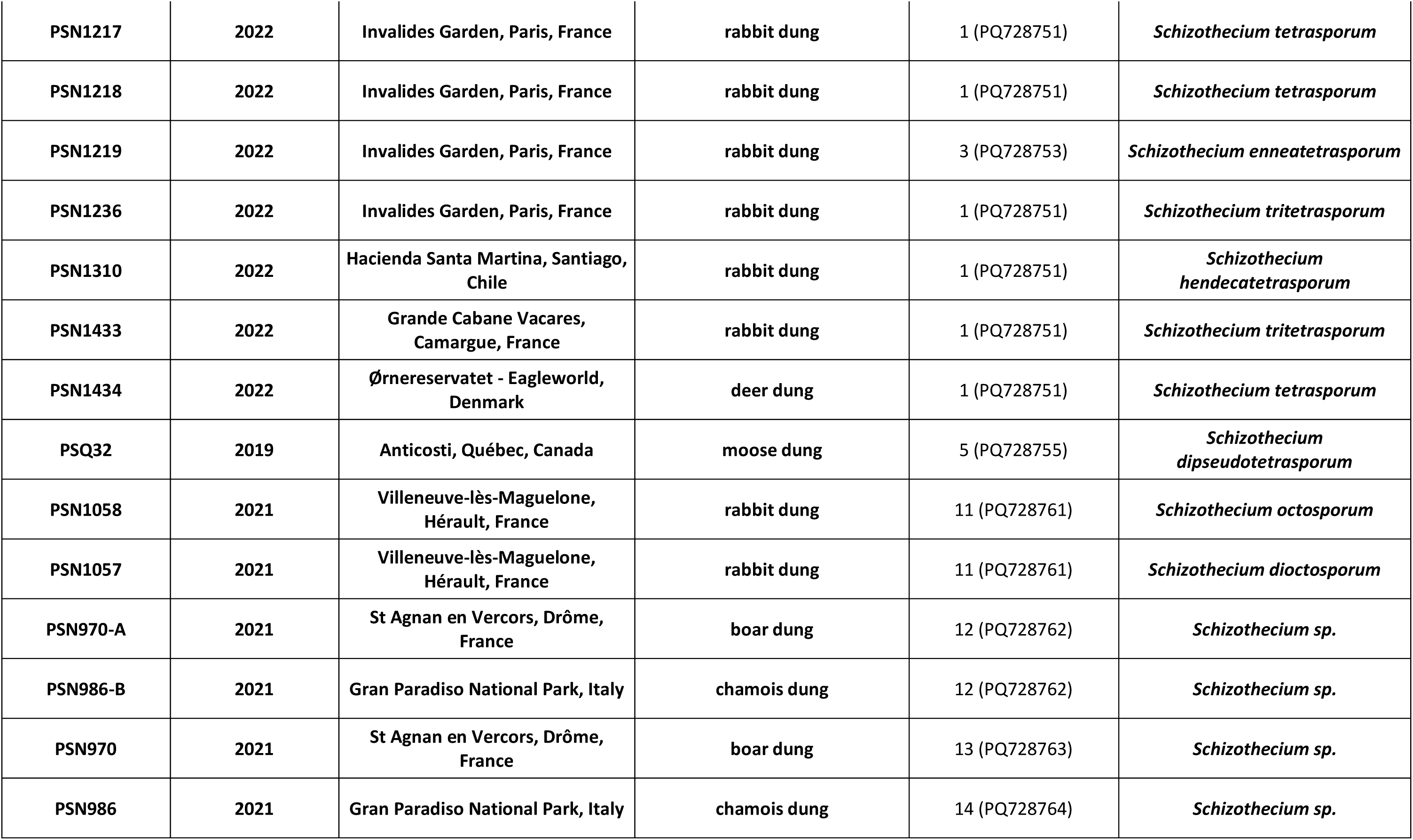
Strains used in this study, isolation date, geographical origin, substrate, ITS sequence type and species.

Dung samples were incubated in moist chambers and strains were isolated as described previously (Silar 2020). Soil samples were incubated and strains were isolated as described in (Silar et al. 2025). All strains were cultivated as described in (Silar 2020) and crossed as in (Vittorelli et al. 2023). The epitype and type specimens were deposited in the Herbarium of the “Museum National d’Histoire Naturelle” (MNHN, Paris, France) and the ex-type living cultures in the “Centre International de Ressources Microbiennes - Champignons Filamenteux” (CIRM-CF, INRAE, France) when these were not already available in collections.

Homokaryons were obtained either by fragmenting heterokaryons or by collecting “small ascospores” from fertile heterokaryons as described in (Silar 2020).

### DNA extraction, ITS and genome sequencing

Rapid DNA extraction for ITS sequencing was performed as described in (Lecellier and Silar 1994) or in (Silar et al. 2025). The ITS region was amplified using the ITS1 and ITS4 primers (White et al. 1990) and the resulting PCR products were sequenced by Genewiz from Azenta (Takeley, UK) using the same primers.

The genome of the CBS815.71 strain has previously been sequenced (Vittorelli et al. 2023). For genome sequencing of the other strains, DNA was extracted using the NucleoSpin® Soil from Macherey Nagel (Düren, Germany) and submitted to 2×150 bp Novaseq illumina sequencing by either Novogene (Cambridge, United Kingdom), Genoscope (Évry-Courcouronnes, France) or Genewiz (Leipzig, Germany). The 82 genomes had an average sequencing coverage of 73X.

### Genome assembly and ANI analysis for species delimitation

The genomes were assembled using Unicycler (Wick et al., 2017) (Table S1). We screened the obtained assemblies for contamination and cleaned the 35 contaminated genomes with the FCS-GX (Foreign Contamination Screen) tool suite from the National Center for Biotechnology Information’s (NCBI) (Astashyn et al. 2024). ANI calculations were made with FungANI (Lalanne and Silar 2024) and the clustering were made with the R statistical software (version 4.3.3) to identify genomes likely belonging to delimit species. Hierarchical clustering using complete linkage method was applied to the distance matrix computed from pairwise Euclidean distance between ANI values.

### Mapping, SNP calling and phylogeny

The ITS phylogenetic tree was constructed by aligning the sequences with MAFFT version 5 (Katoh et al. 2005) and by producing the phylogenetic tree using PhyML with the default parameters (Guindon and Gascuel 2003). The tree was visualized using FigTree (Rambaut 2007).

We mapped the 82 Illumina genomes, including CBS815.71, to the CBS815.71sp3 genome assembly (Vittorelli et al. 2023) and performed SNP calling in order to produce a phylogeny based on SNPs. Read trimming, mapping and SNP calling were performed using the pipeline described previously (Hartmann et al. 2021). We only kept good-quality and biallelic SNPs called in at least 90% of our strains. To build a phylogeny of the 82 strains, we identified SNPs located in BUSCO genes (Manni et al. 2021) from the *Sordariomycetes* ortholog set (sordariomycetes_odb10, 2020-08-05) using bedtools v2.26.0 (Quinlan and Hall 2010) and selected them using vcftools v0.1.17 (Danecek et al. 2011). We converted the obtained file from vcf to phylip with PGDSPider v.2.1.1.5 (Lischer and Excoffier 2012). We computed a maximum-likelihood based phylogeny with IQ-TREE v1.6.1 (Nguyen et al. 2014; Kalyaanamoorthy et al. 2017), using ModelFinder to find the best fit model and ultrafast bootstraps of 1000 iterations with the following options: -s -bb 1000 -alrt 1000. Our final dataset for this phylogeny contained 1,166,408 SNPs. We manually rooted the phylogeny with the ape package in R (Paradis and Schliep 2018), using the strains PSN970 and PSN986 as the outgroup. These species were the most divergent of the sequenced set based on an ITS barcode-based phylogeny including sequences from the other *Schizothecium* species available in GenBank RefSeq (data will be presented in a forthcoming paper on the genus *Schizothecium* diversity). We represented the phylogeny with the ggtree package in R (Yu 2020; Yu et al. 2018; Yu et al. 2017).

### Morphological analyses

All species differentiated perithecia on V8 medium at 22°C and expelled ascospores after two to three weeks of incubation, as previously described for the CBS815.71 strain (Vittorelli et al. 2023). Perithecia of the eight-spored species were obtained on V8 medium after three to four weeks. All four-spored and eight-spored strains were also fertile on M2, oat meal agar and M0 medium supplemented with shredded miscanthus. However, perithecium maturation was delayed one or two weeks on these media and produced in fewer amounts, especially on M2. For whole perithecium illustrations, pictures were taken from M0 + miscanthus plates as perithecia were there more scattered and more easily visualized than on other media. Perithecium analyses were made on fruiting bodies obtained on V8 and starting to expel ascospores. Ascospores for size measurements were collected on cover agar plates (Silar 2020). The perithecium and primary appendage sizes were measured on 10 different perithecia and ascospores, respectively, and ascospore heads sizes on 50 different dikaryotic ascospores.

Of note, many isolates of the four-spored species often lost the nuclei of one mating type when cultivated in Petri plates for a long time, as previously described for strain CBS815.71 (Vittorelli et al. 2023), rendering such cultures barren. Additionally, the eight-spored PSN1057 and PSN1058 strains frequently lost fertility following culture, although no alteration of mycelium morphology was observed.

## Results and Discussion

### Isolation of four-spored strains

To assess the diversity of *S. tetrasporum*, we obtained 76 four-spored strains with typical *S. tetrasporum* morphology either from culture collections or directly from nature (Table 1). Newly isolated strains were mostly from dung, especially those of rabbits and hares, in line with the known biology of *S. tetrasporum* (see (Lundqvist 1972) page 256 and (Bell and Mahoney 1995)), but some were from soil or mud, indicating a possible broader biotope than considered for these species. The recovered strains were from various origins, mostly from Western Europe, especially from France (56 isolates; Fig. 1).

**Figure 1.**
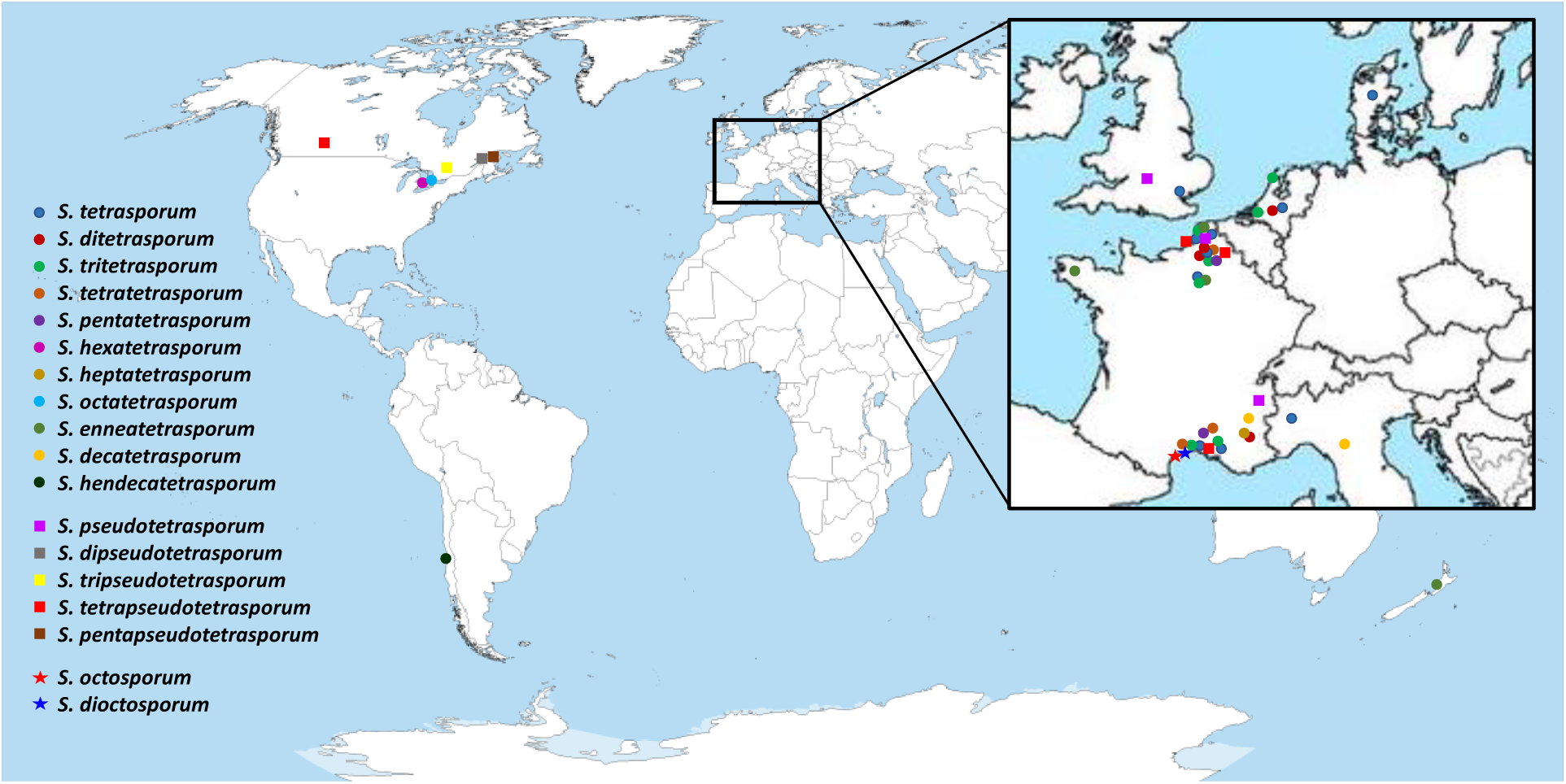
Isolate origins.

### ITS and genome sequencing of four-spored and eight-spored strains

As a first step, we sequenced the ITS region of the four-spored strains and found ten different ITS sequences (Table 1, Fig. 2), highly similar to those of *Schizothecium spp.*: they displayed 99-100% identity with the ITS of the previously sequenced CBS815.71 genome and 97-98% identity with the ITS of “*Schizothecium vesticola* strain SMH3187-1” sequenced by the JGI (Hensen et al. 2023). Such diverse ITS sequences and morphological similarity suggested the existence of different sibling species. To test this hypothesis, we sequenced their genomes with the Illumina technology, along those of six strains of *Schizothecium spp.* having eight-spored asci (see Table 1 for the origins of these strains, which may be found at the same locations at the four-spored strains) and chosen because they also had ITS sequences highly similar to those of the four-spored ones (Table1, Fig. 2; 98-99% identity with those of the four-spored strains). Interestingly, two of these strains (PSN1057 and PSN1058 with the ITS sequence #11) had their ITS sequences nested within those of the four-spored ones in a PhyML tree (Fig. 2).All four-spored strains had a genome size comprised between 28.5 and 32.7 Mb (see Table S1 for genome assembly statistics). The sequences of the 76 *S. tetrasporum* (*sensu lato*) strains were analyzed with FungANI (Lalanne and Silar 2024) followed by clustering analyses of the ANIs. The data showed that the 76 *S. tetrasporum s. l.* strains actually belonged to 16 distinct species with intraspecies ANI higher than 99.5% (Fig. 3, see supplementary data 1 for all ANI results). The 16 species were distributed into two distinct clusters, with ANI values for strains within clusters ranging from 97.5 to 99.2% (for PSN1042/CBS370.79 and PSN663/PSN589, respectively) and ranging from 90.2 to 91.9% for strains belonging to different ones (PSN377/PSN1042 and PSN668/PSN830, respectively).

**Figure 2.**
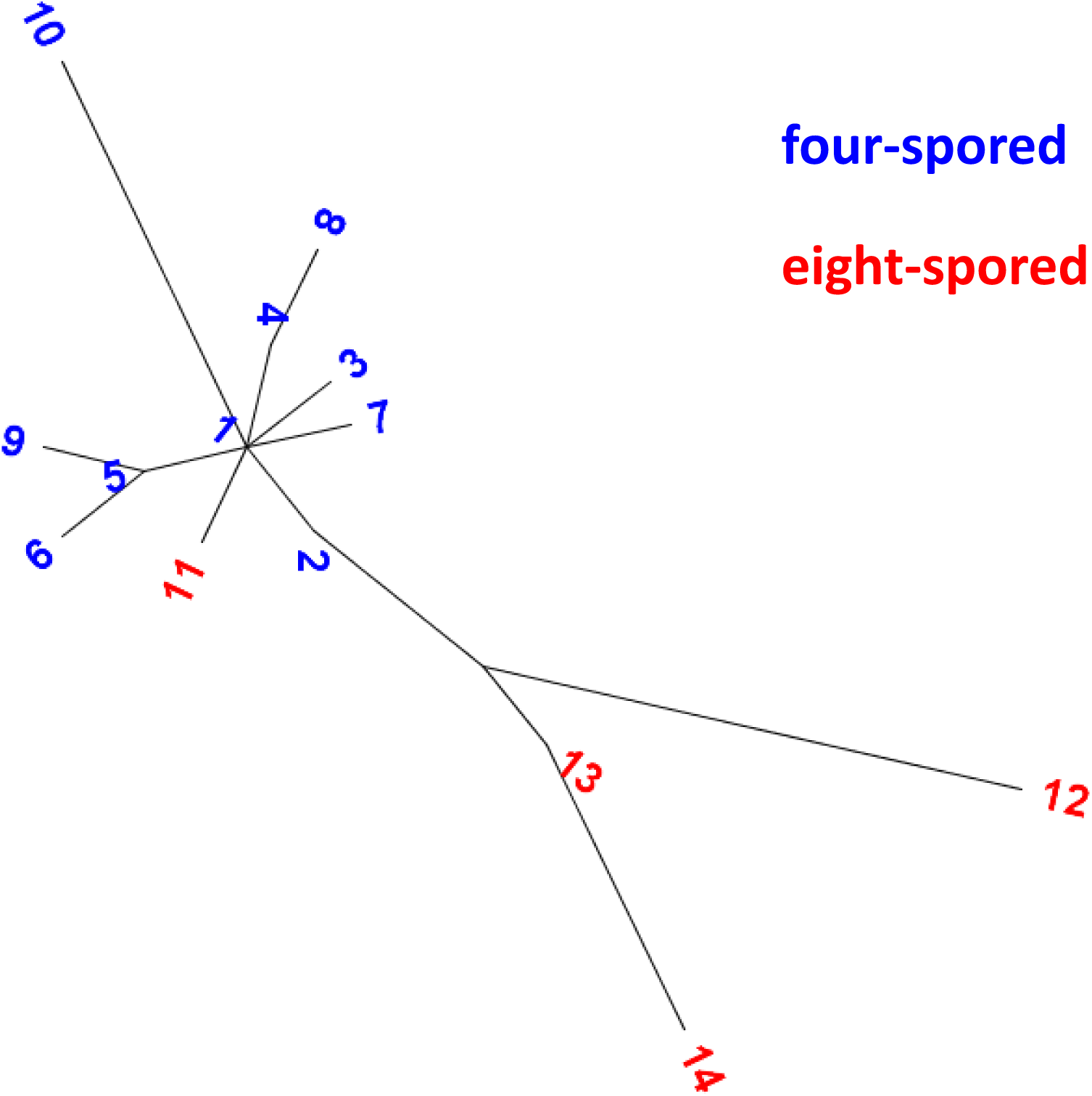
PhyML tree of the 10 different ITS sequences of four spored strains (in blue) along with those of eight-spored ones (in red). The numbers at the tip are the ITS sequence ID used in Table 1.

**Figure 3.**
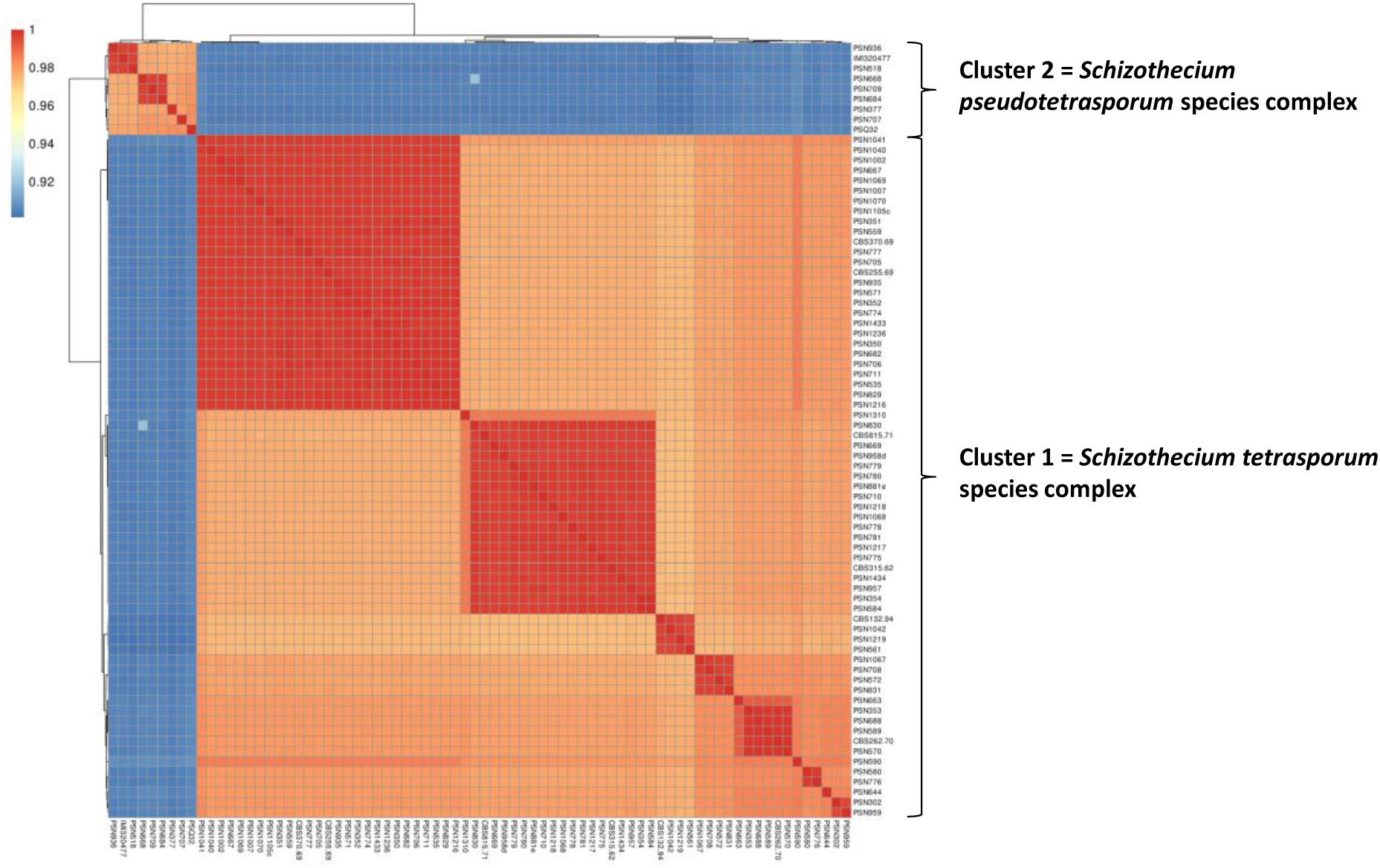
Clustering analysis of the ANI (average nucleotide identity) of four-spored strains.

The numbers of generations necessary to obtain 1‰, 1% and 3.7% sequence divergence were estimated in *Saccharomyces cerevisiae* to be ∼2 million, ∼20 million and ∼60 million, respectively, using the known mutation rate for this species of about 3 x 10^-10^ mutations per generation and nucleotide (Rolland and Dujon 2011). These ages are likely underestimated as all mutations were assumed neutral and independent, while the elimination of deleterious mutations by natural selection should slow down mutation accumulation. The divergence between two *Colletotrichum* species (*C. incanum* H.-C. Yang, J.S. Haudenshield & G.L. Hartman and *C. tofieldiae* (Pat.) Damm, P.F. Cannon & Crous) with an ANI of 89% was estimated to have occurred 8.8 million year ago (Hacquard et al. 2016). We estimated the possible divergence times within the *S. tetrasporum* complex assuming a mutation rate similar to that of *S. cerevisiae* and five generations per year. Indeed, at most 15 generations/year could be obtained in optimal laboratory conditions never met in the wild, for example because temperature in most of autumn, winter and spring would be too low for growth. We thus estimated that the two species clusters have diverged 10 to 15 million years ago, and species from within cluster about 5 million years ago. These time values estimated using Rolland and Dujon’s method are thus similar to the value found by Hacquard et al. for fungi having an ANI of about 90%. As a point of reference, 10-15 million years and 5 million years roughly correspond to the divergence of the gorilla and chimpanzee lineages from the human one, respectively (Langergraber et al. 2012). The ANI between the human and gorilla genomes is 98.3 (Scally et al. 2012) and that of between human and chimpanzee is 98.6% (Suntsova and Buzdin 2020). Note that calculations are somewhat different in the cases of apes/Humans because although Humans and apes have longer generation times (20-30 years), they likely have a higher mutation rate (1 x 10^-9^ mutation per generation and nucleotide; see (Langergraber et al. 2012)). Hence, the diversity found here in the *S. tetrasporum* complex likely reflects ancient divergence events.

One of the two clusters was more species-rich than the other (eleven species versus five), with also more strains per species (up to 27 isolates for one species). Noteworthy, several species were represented by a single strain, suggesting that many more species are likely present in nature. The most common ITS sequence was shared by eight species. One species (hereafter named *Schizothecium tritetrasporum*) had three different ITS sequences. The most frequent ITS sequence (#1) was shared by species from the two ANI-defined species clusters (Table 1).

The genomes of the eight-spored strains were slightly larger than those of the four-spored ones, ranging from 33.7 Mb to 36.9 Mb. FungANI analyzes showed that the eight-spored strains belonged to six different species (with ANI ranging from 87.0% to 95.7%, see supplementary data 2) and that they likely clustered into three species complexes (Fig. 4). Of interest, PSN1057 and PSN1058 were assigned to different species despite having the same ITS; this was also the case for PSN970 and PSN986. Note that strains PSN970, PSN970-A, PSN986 and PSN986-B were not investigated in depth for the present paper and will be described in a forthcoming paper on the whole *Schizothecium* genus diversity.

**Figure 4.**
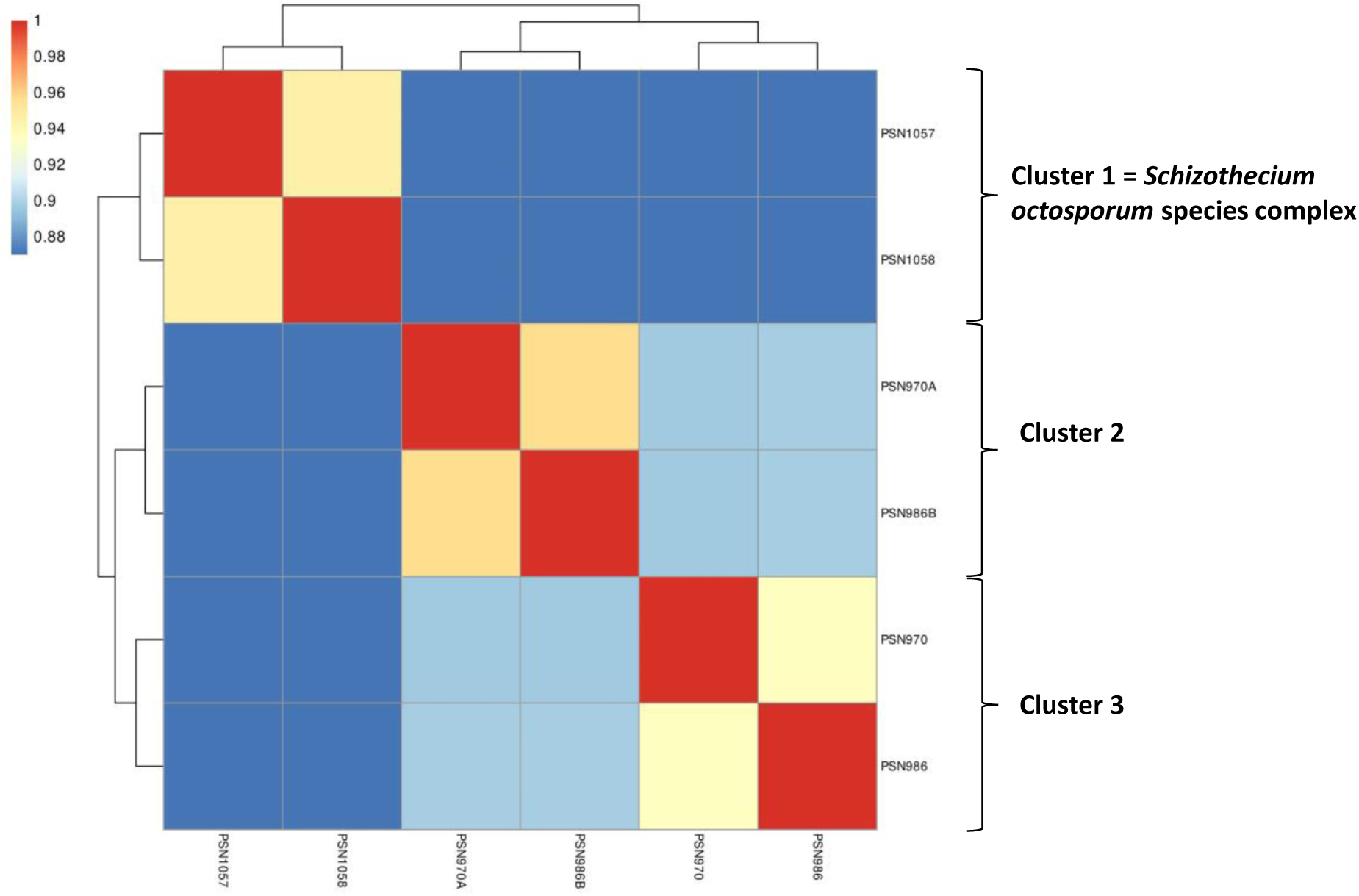
Clustering analysis of the ANI of eight-spored strains.

### The phylogenetic tree based on SNPs confirms the ANI analysis

To better characterize the relationships among the different species, we built a phylogenetic tree based on the SNPs of the 81 new genomes together with that of the previously sequenced strain CBS815.71 (Vittorelli et al. 2023). We rooted the tree with the two most-distant eight-spored strain PSN970 and PSN986 (Fig. 5). Analysis of the phylogenetic tree based on genome-wide SNPs confirmed with high statistical support the existence of the two distinct and distantly-related clusters of four-spored strains defined by the ANI analysis. Interestingly. the tree also confirmed with high statistical support the placement between these two clusters of the two eight-spored strains with an ITS branching within the four-spored strain ones. Therefore, pseudo-homothallism in *S. tetrasporum sensus lato* may have evolved twice independently or evolved once ancestrally in the clade, with a reversion to heterothallism in the ancestors of the three eight-spored strains.

**Figure 5.**
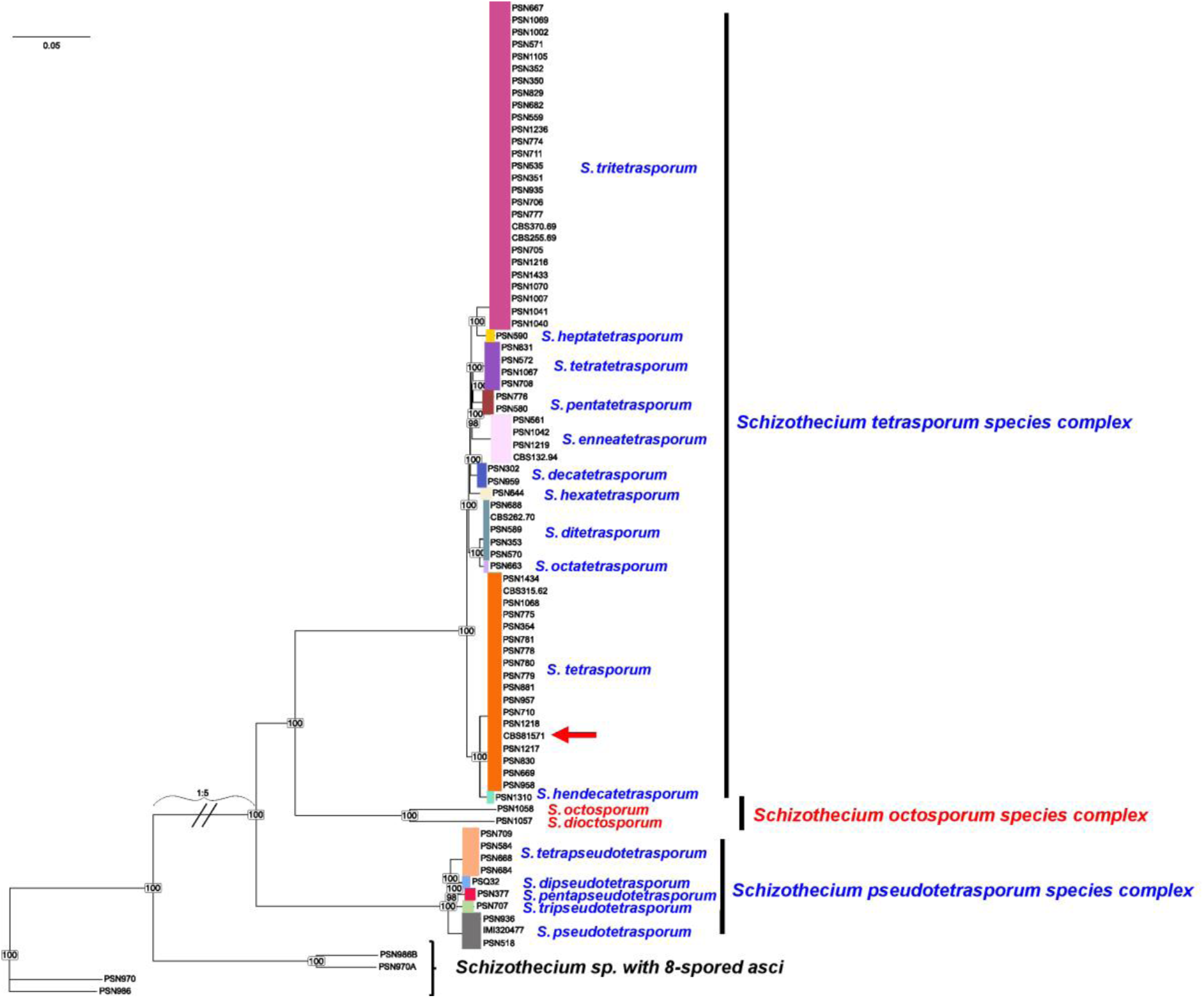
Phylogenomic tree based on genome-wide SNPs with strain CBS815.71 (red arrow). Numbers at nodes indicates bootstrap values of 1000 iterations. Colored vertical rectangles delimit species.

We chose to define three species complexes, corresponding to the three clades separated from each other by long branches, and with contrasting breeding systems (four-versus eight-spored asci). The first complex contained the previously sequenced strain CBS815.71, which we defined as ex-epitype for S*. tetrasporum sensus stricto* and which was chosen as the representative for the *S. tetrasporum* species complex. The second species complex contained the available commercial strain IMI320477. We named it the *S. pseudotetrasporum* species complex. We do not propose strain IMI320477 as representative for this complex because this strain had nearly colorless ascospores, likely because of a mutation. Instead, we propose strain PSN936 as the type strain for *S. pseudotetrasporum* and representative for this complex. The third one contained the two eight-spored strains PSN1057 and PSN1058 and was therefore named the *S. octosporum* species complex. The PSN1058 strain was chosen as type for the *S. octosporum* species and as a representative of this complex. Morphological analyses showed that species from this complex did not fit with any previously known species (see Fig. 22 and 23). We decided to name the different species in each cluster by following a rule similar to the one used to name the 14 sibling species of the *Paramecium aurelia* complex (Sonneborn 1975). We added a numeral prefix derived from Greek to the name of the complex, but we left the species type for each complex without such a prefix. Although this rendered the naming a little bit complex and sometime awkward (such as in “*tetratetrasporum*”), this naming rule allows to easily attribute to which complex each species belonged and should provide a convenient rule for the naming of other species discovered in the future. Note that we expect such species to be numerous and having a rule would avoid a plethora of names based on discovery location, substrate or person names that may not be representative of the morphology of the species.

The two species of the *S. octosporum* complex were found at the same small area in the south of France (Table 1) and only there. On the contrary, both four-spored species complex seemed to have a large range distribution. For example, *S. tetrapseudotetrasporum* was found in Canada, as well as in the south and the north of France (Table 1, Figure 1). Some species appeared less cosmopolitan, such as *S. tetrasporum* and *S. tritetrasporum*, for which the numerous isolates were all from western Europe. However, only few locations were sampled, so that inference on distribution ranges is premature and the diversity revealed in the present study likely only represents a small fraction of the actual one. For example, analysis of a limited number of samples from Chile identified a species not present in other sampled locations, and samples from Canada revealed six different species, including five only found there so far (Table 1, Figure 1). Multiple sibling species co-existed in small areas (Table 1), such as in the Invalides Garden in Paris, famous for its rabbit population, where we detected three different species. Different species could even be isolated from the same dung batch, as for example PSN589 and PSN590 (belonging to *S. ditetrasporum* and *S. heptatetrasporum*, respectively), which were isolated from the same batch of hare dung. Although we analyzed samples coming from tropical regions, we have not found any *Schizothecium* fungus there, suggesting that these fungi, including *S. tetrasporum*, are restricted to cold climates, in line with our observations that *Schizothecium* species would not fructify in the laboratory at temperatures above 22/23°C.

### Comparative morphological analysis of the four-spored species

We then performed comparative morphological analyses on ex-type strains to assess whether the different species could be differentiated based on the morphology of their sexual reproduction structures or their mycelium growth speed. As seen in Figs. 6-21 and Table 2, small interspecies differences could be observed for the four-spored species. To test if these differences were higher than intraspecific variation, we analyzed seven strains of *S. tetrasporum s.s.* (*sensu stricto*) in addition to CBS815.71 (PSN669, PSN710, PSN781, PSN830, PSN957, PSN958, PSN1068, Table 3). As seen in Table 2, intraspecific differences within *S. tetrasporum s.s.* were similar to those between the different four-spored species, indicating that the different four-spored species cannot be distinguished on the basis of morphology of the sexual reproduction structures or growth speed with certainty. Moreover, we did not observe differences between the sibling species in their repartition of fruiting bodies on the thallus on any of the media tested, unlike for the *P. anserina* species complex (Boucher et al. 2017). nevertheless, the different species discovered here were not studied as much as the *P. anserina* species. Noteworthy, we noticed that fruiting body production was much more variable than in *P. anserina* because most species seemed to easily loose nuclei from one mating type, as previously shown for strain CBS815.71 (Vittorelli et al. 2023).

**Fig. 6.**
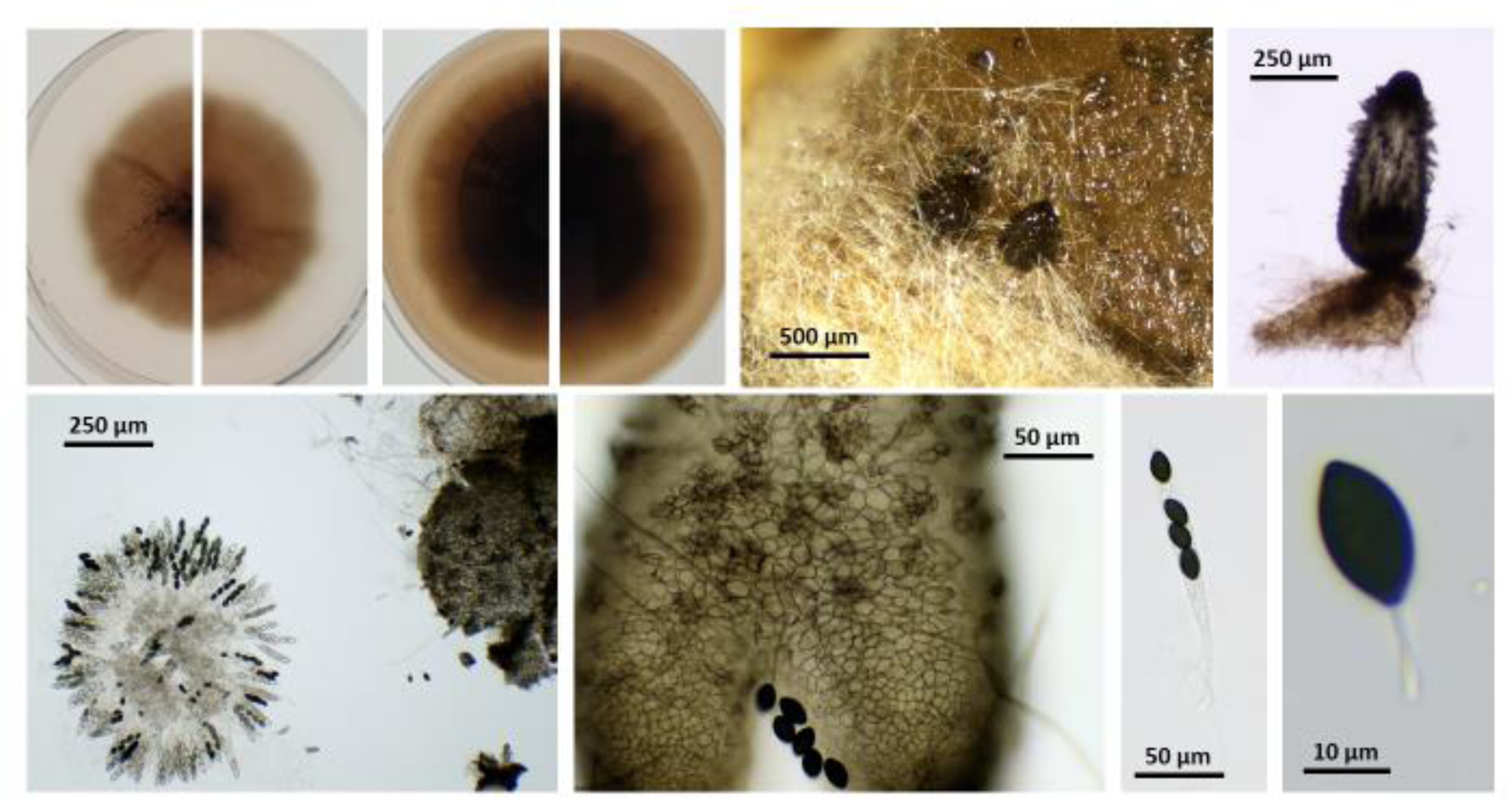
CBS815.71, the ex-epitype of *Schizothecium tetrasporum*. Top from left to right: culture on M2 top view, culture on M2 reverse view, culture on V8 top view, culture on V8 reverse view, perithecia on M0+ miscanthus, isolated perithecia. Bottom from left to right; squeezed perithecium with rosette of asci and some peridium, enlarge peridium view, ascus and ascospore.

**Table 2.** Morphological features of selected four-spored and eight-spored strains of the *Schizothecium tetrasporum* species complexes. Mycelium growth speed on V8 and M2, sizes of perithecium, primary appendage and ascospore heads (means and standard deviations), and percentage of homokaryotic ascospores.

|  |  | Mycelium growth speed<br>mm/day |  |  | ascospores size |  | % of small<br>homokaryotic<br>ascospores |
| --- | --- | --- | --- | --- | --- | --- | --- |
| Species | ex-type | on V8 | on M2 | perithecium size<br>(height x width μm) | spore head (μm) | primary appendage (μm) |  |
| S. tetrasporum complex |  |  |  |  |  |  |  |
| S. tetrasporum | CBS815.71 | 1.4 +/- 0.0 | 1.1+/- 0.1 | 630 +/- 50 x 230 +/- 20 | 23.6 +/- 1.8 x 13.6 +/- 0.7 | 14.9 +/- 2.4 x 2.3 +/- 0.4 | 5.90% |
|  | PSN669 | 1.2 +/- 0.1 | 1.1 +/- 0.1 | 440 +/- 50 x 220 +/- 20 | 24.9 +/- 1.3 x 14.3 +/- 0.9 | 11.4+/- 2.0 x 3.0 +/- 0.5 | 2.80% |
|  | PSN710 | 1.2 +/- 0.0 | 1.1 +/- 0.1 | 370 +/- 30 x 200 +/- 20 | 25.5 +/- 1.6 x 13.9 +/- 1.0 | 11.2 +/- 2.2 x 3.2 +/- 0.9 | 7.00% |
|  | PSN781 | 1.3 +/- 0.1 | 1.0 +/- 0.1 | 540 +/- 30 x 250 +/- 30 | 26.2 +/- 1.3 x 15.1 +/- 0.6 | 12.1 +/- 2.3 x 3.1 +/- 0.9 | 0.90% |
|  | PSN830 | 1.4 +/- 0.0 | 1.1 +/- 0.1 | 450 +/- 60 x 240 +/- 30 | 25.0 +/- 1.4 x 13.3 +/- 0.7 | 9.3 +/- 1.3 x 2.2 +/- 0.4 | 5.30% |
|  | PSN957 | 1.3 +/- 0.0 | 1.0 +/- 0.1 | 390 +/- 50 x 200 +/- 20 | 25.9 +/- 1.4 x 14.2 +/- 0.8 | 11.7 +/- 1.2 x 2.5 +/- 0.4 | 9.50% |
|  | PSN958 | 1.3 +/- 0.0 | 1.1 +/- 0.1 | 540 +/- 70 x 250 +/- 30 | 25.5 +/- 1.4 x 14.5 +/- 0.6 | 14.2 +/- 2.3 x 2.7 +/- 0.5 | 0.90% |
|  | PSN1068 | 1.3 +/- 0.1 | 1.2 +/- 0.1 | 490 +/- 40 x 240 +/- 20 | 24.6 +/- 1.2 x 14.3 +/- 0.6 | 8.5 +/- 1.4 x 2.1 +/- 0.3 | 13.60% |
| S. ditetrasporum | PSN589 | 1.3 +/- 0.0 | 1.1+/- 0.1 | 490 +/- 70 x 270 +/- 20 | 25.1 +/- 1.4 x 14.6 +/- 0.6 | 13.2 +/- 1.8 x 2.6 +/- 0.5 | 1.50% |
| S. tritetrasporum | CBS370.69 | 1.4 +/- 0.0 | 1.2 +/- 0.1 | 490 +/- 50 x 230 +/- 20 | 27.0 +/- 1.6 x 14.8 +/- 0.6 | 10.5 +/- 1.0 x 3.0 +/- 0.7 | 0.25% |
| S. tetratetrasporum | PSN831 | 1.4 +/- 0.0 | 1.0 +/- 0.1 | 360 +/- 40 x 220 +/- 20 | 24.5 +/- 1.4 x 13.3 +/- 0.7 | 14.5 +/- 1.4 x 3.0 +/- 0.6 | 2.80% |
| S. pentatetrasporum | PSN580 | 1.4 +/- 0.0 | 1.0 +/- 0.1 | 450 +/- 50 x 280 +/- 30 | 26.3 +/- 1.2 x 14.2 +/- 0.7 | 11.5 +/- 0.6 x 2.7 +/- 0.5 | 2.30% |
| S. hexatetrasporum | PSN644 | 1.4 +/- 0.1 | 1.1+/- 0.1 | 440 +/- 40 x 230 +/- 20 | 24.1 +/- 1.2 x 14.5 +/- 0.9 | 10.9 +/- 1.1 x 2.7 +/- 0.4 | 2.10% |
| S. heptatetrasporum | PSN590 | 1.2 +/- 0.0 | 1.0 +/- 0.1 | 460 +/- 40 x 230 +/- 20 | 25.6 +/- 2.0 x 15.3 +/- 1.2 | 12.0 +/- 1.9 x 2.4 +/- 0.5 | 0.33% |
| S. octatetrasporum | PSN663 | 1.5 +/- 0.1 | 1.3 +/- 0.1 | 510 +/- 44 x 250 +/- 10 | 26.4 +/- 1.4 x 15.0 +/- 0.4 | 11.1+/- 1.4 x 2.4 +/- 0.3 | 2.00% |
| S. enneatetrasporum | PSN1042 | 1.4 +/- 0.1 | 1.1 +/- 0.0 | 520 +/- 50 x 230 +/- 20 | 26.4 +/- 1.0 x 15.2 +/- 0.5 | 11.2 +/- 1.8 x 2.7 +/- 0.4 | <0.1% |
| S. decatetrasporum | PSN959 | 1.1 +/- 0.1 | 1.1 +/- 0.0 | 430 +/- 30 x 210 +/- 10 | 26.3 +/- 1.1 x 13.6 +/- 0.7 | 12.5 +/- 1.7 x 2.6 +/- 0.5 | 2.25% |
| S. hendecatetrasporum | PSN1310 | 1.4 +/- 0.1 | 1.0 +/- 0.1 | 450 +/- 41 x 210 +/- 10 | 26.9 +/- 0.9 x 15.3 +/- 0.6 | 13.0 +/- 1.5 x 3.2 +/- 0.6 | 1.00% |
| S. pseudotetrasporum complex |  |  |  |  |  |  |  |
| S. pseudotetrasporum | PSN936 | 1.4 +/- 0.1 | 1.1 +/- 0.1 | 490 +/- 40 x 260 +/- 30 | 26.0 +/- 1.1 x 14.4 +/- 0.5 | 8.8 +/- 1.4 x 2.6 +/- 0.2 | 1.80% |
| S. dipseudotetrasporum | PSQ32 | 1.4 +/- 0.1 | 1.1 +/- 0.1 | 370 +/- 40 x 200 +/- 10 | 24.4 +/- 1.5 x 13.9 +/- 0.5 | 10.5 +/- 1.4 x 2.9 +/- 0.3 | 3.70% |
| S. tripseudotetrasporum | PSN707 | 1.4 +/- 0.0 | 1.3 +/- 0.1 | 450 +/- 40 x 234 +/- 30 | 25.0 +/- 1.0 x 13.8 +/- 0.6 | 7.8 +/- 1.1 x 2.4 +/- 0.5 | 1.30% |
| S. tetrapseudotetrasporum | PSN684 | 1.4 +/- 0.1 | 1.1 +/- 0.0 | 400 +/- 50 x 250 +/- 30 | 26.4 +/- 1.4 x 14.2 +/- 0.6 | 13.8 +/- 1.3 x 2.7 +/- 0.3 | 1.00% |
| S. pentapseudotetrasporum | PSN377 | 1.6 +/- 0.1 | 1.2 +/- 0.1 | 480 +/- 40 x 270 +/- 20 | 26.9 +/- 2.1 x 14.8 +/- 1.1 | 10.2 +/- 1.5 x 2.3 +/- 0.4 | 0.90% |
| S. octosporum complex |  |  |  |  |  |  | % of large ascospores |
| S. octosporum | PSN1058 | 2.3 +/- 0.1 | 1.6 +/- 0.3 | 600 +/- 50 x 290 +/- 40 | 19.1 +/- 1.5 x 11.0 +/- 0.8 | 10.1 +/- 0.4 x 1.7 +/- 0.4 | NA |
| S. dioctosporum | PSN1057 | 2.3 +/- 0.0 | 1.6 +/- 0.1 | 500 +/- 60 x 380 +/- 60 | 19.0 +/- 0.3 x 11.0 +/- 0.1 | 7.9 +/- 1.5 x 1.8 +/- 0.3 | NA |

### Comparative morphological analysis of the eight-spored strains PSN1058 and PSN1057

Morphological analyses of PSN1058 and PSN1057, the two heterothallic strains nested within the pseudo-homothallic ones revealed a clear difference in perithecium morphology. Indeed, while PSN1058 produced slender perithecia looking very much like those of the four-spored species (Fig. 22), the perithecia of PSN1057 were globose, being almost as wide as tall (Fig. 23). Both strains produced slightly asymmetrical ascospores, but not as much as *Schizothecium fimbriatum* (A. Bayer) Barrasa & Soláns or *Schizothecium vratislaviense* (Alf. Schmidt) Doveri & Coué. ITS sequences indicate that PSN1058 and PSN1057 are closely related to *Schizothecium vesticola* (Berk. & Broome) N. Lundq. Strains from this species produce eight-spored asci and are likely heterothallic (Bell and Mahoney 1995). However, this species has symmetrical ascospores.

Finally, to confirm species delimitation, we performed intra-specific crosses between five strains of *S. tetrasporum s. s.* (CBS815.71, PSN781, PSN957, PSN710 and PSN1068; 10 different cross combinations) and five strains of *S. tritetrasporum* (CBS370.69, PSN571, PSN535, PSN352 and PSN935; 10 different cross combinations), as well as inter-species crosses between the two chosen species (50 different cross combinations). All interspecies crosses were sterile, while one and three intraspecific crosses were fertile, for *S. tetrasporum s.s.* and *S. tritetrasporum*, respectively. Ascospores collected from the intraspecific crosses germinated as well as those from intra-strain crosses. The lack of inter-species fertile crosses reinforced the view that these two species defined by genomic criteria were *bona fide* species. Note that lack of success of crosses within the same species has previously been documented in *Sordariales* and may be due to heterokaryons incompatibility (Ament-Velásquez et al. 2022).

### Taxonomy

***Schizothecium tetrasporum*** (Wint.) Lundqv., in Symb. Bot. Upsal. 20(no. 1): 256 (1972). Index Fungorum number: 323145; Fig. 6

*Synonyms*: *Sordaria tetraspora* G. Winter, in Hedwigia 11: 161 (1872) – IF=225188; *Podospora tetraspora* (G. Winter) Cain, in Can. J. Bot. 40: 460 (1962) – IF=337430; *Neoschizothecium tetrasporum* (G. Winter) S.K. Huang & K.D. Hyde, in Huang, Hyde, Mapook, Maharachchikumbura, Bhat, McKenzie, Jeewon & Wen, Fungal Diversity: 10.1007/s13225-021-00488-4, [98] (2021) -IF=558342

Typus: Feb. 1872, herb. Barbey-Boissier 1529 (NEOTYPE, G) on mouse dung from Harth bei Leipzig, see (Bell and Mahoney 1995), epitype (PC0798995,designated here); ex-epitype : CBS815.71

Description: mycelium slow growing (1.1 +/- 0.1 mm/day on M2, 1.4 +/- 0.0 mm day on V8), dark green to almost black with few aerial hyphae on M2 and profuse ones on V8, often but not always undergoing lysis. Perithecia conical to ovoid, 630 +/- 50 x 230 +/- 20 µm, membranous, pale brown, semi-transparent when young, darker to almost black when old. Neck not-clearly differentiated, sometimes slightly curved towards the light, decorated by small triangular short swollen hairs typical of *Schizothecium* perithecia. Peridium with *textura angularis*. Asci four-spored and claviform. Spores obliquely uniseriate, transversely septate with hyaline primary appendage and secondary appendages at both poles often difficult to see. Spore head 23.6 +/- 1.8 x 13.6 +/- 0.7 µm, ellipsoidal, smooth, flattened at the base, with an apical germ pore. Primary appendage (pedicel) cylindric, slightly tapering towards the apex, 14.9 +/- 2.4 x 2.3 +/- 0.4 µm. Secondary appendage present but evanescent and lacking on most expelled ascospores.

Habitat & Distribution: frequent on dung, especially rabbit ones. Possibly present also in soil (epitype CBS815.71 was isolated from forest soil in the U.K.). Isolated from Western Europe (France, Italy, United-Kingdom, Netherlands and Denmark).

Mating strategy: pseudo-homothallic.

Additional strains : Denmark, PSN1434. France, PSN354, PSN669, PSN775, PSN778, PSN779, PSN780, PSN781, PSN830, PSN881, PSN957, PSN958, PSN1068, PSN1217, PSN1218. Italy, PSN710. Netherlands, CBS315.62.

***Schizothecium ditetrasporum*** Silar ***sp. nov.*** Fig. 7

**Fig. 7.**
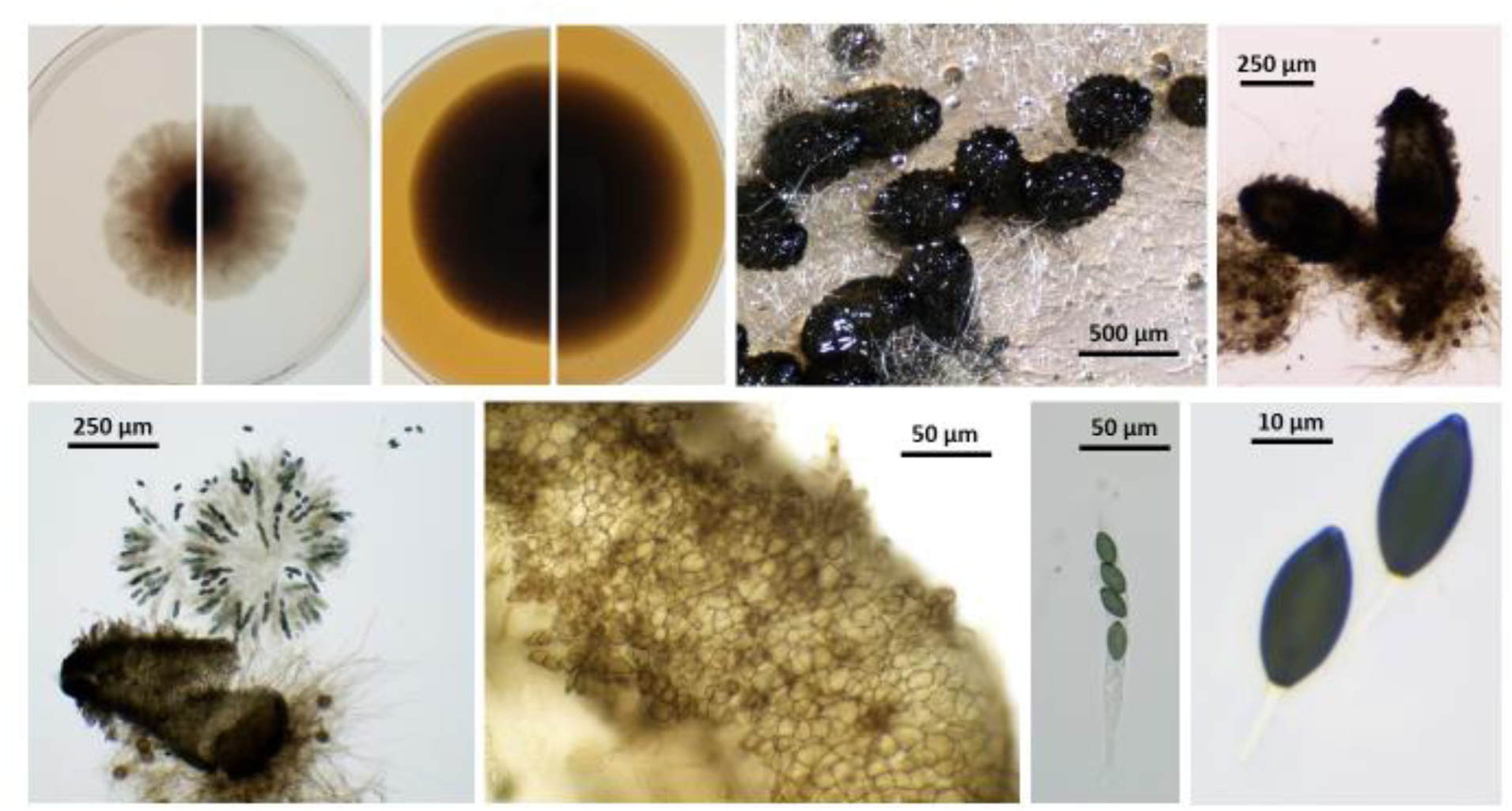
PSN589, the ex-type of *Schizothecium ditetrasporum*. Top from left to right: culture on M2 top view, culture on M2 reverse view, culture on V8 top view, culture on V8 reverse view, perithecia on M0+ miscanthus, isolated perithecia. Bottom from left to right; squeezed perithecium with rosette of asci and some peridium, enlarge peridium view, ascus and two ascospores.

Index Fungorum number:

*Holotype*: PC0798996, ex-type: PSN589

Description: mycelium slow growing (1.1 +/- 0.1 mm/day on M2, 1.3 +/- 0.0 mm day on V8), dark green to almost black with few aerial hyphae on M2 and profuse ones on V8 and often but not always undergoing lysis. Perithecia conical to ovoid, 490 +/- 70 x 270 +/- 20 µm, membranous, pale brown, semi-transparent when young, darker to almost black when old. Neck not-clearly differentiated, sometime slightly curved towards the light, decorated by small triangular short swollen hairs typical of *Schizothecium* perithecia. Peridium with *textura angularis*. Asci four-spored and claviform. Spores obliquely uniseriate, transversely septate with hyaline primary appendage and secondary appendages at both poles often difficult to see. Spore head 25.1 +/- 1.4 x 14.6 +/- 0.6 µm, ellipsoidal, smooth, flattened at the base, with an apical germ pore. Primary appendage (pedicel) cylindric, slightly tapering towards the apex, 13.2 +/- 1.8 x 2.6 +/- 0.5 µm. Secondary appendage present but evanescent and lacking on most expelled ascospores.

Habitat & Distribution: Saprobic on dung and also found in soil. Isolated from France and the Netherlands. The type is from France (Alpes de Haute Provence) on hare dung.

Mating strategy: pseudo-homothallic.

Additional strains : France: PSN353, PSN570, PSN688. Netherlands: CBS262.70.

***Schizothecium tritetrasporum*** Silar ***sp. nov.*** Fig. 8

**Fig. 8.**
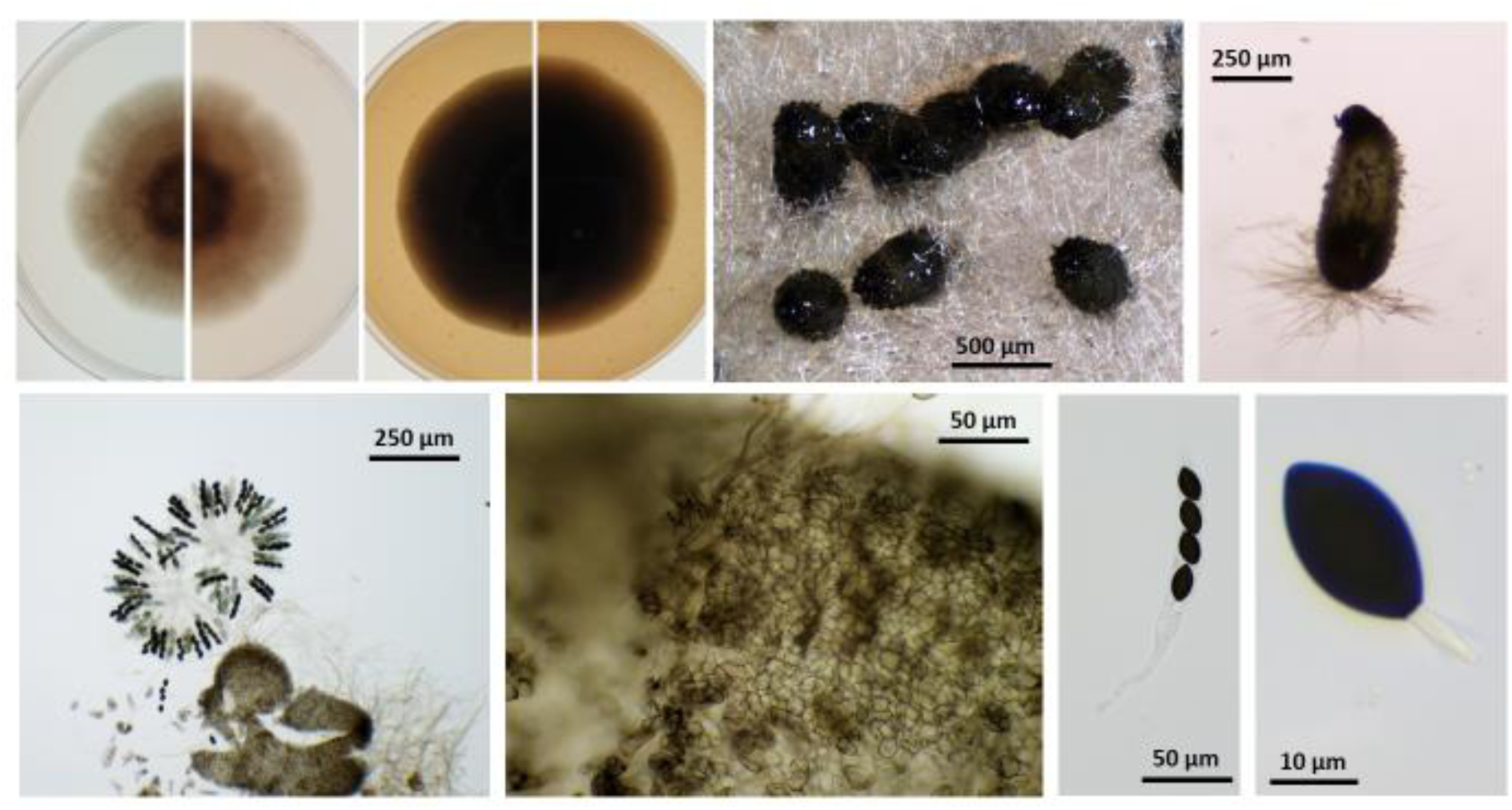
CBS370.69, the ex-type of *Schizothecium tritetrasporum*. Top from left to right: culture on M2 top view, culture on M2 reverse view, culture on V8 top view, culture on V8 reverse view, perithecia on M0+ miscanthus, isolated perithecia. Bottom from left to right; squeezed perithecium with rosette of asci and some peridium, enlarge peridium view, ascus and ascospore.

Index Fungorum number:

*Holotype*: PC0798998, ex-type: CBS370.79

Description: mycelium slow growing (1.2 +/- 0.1 mm/day on M2, 1.4 +/- 0.0 mm day on V8), dark green to almost black with few aerial hyphae on M2 and profuse ones on V8 and often but not always undergoing lysis. Perithecia conical to ovoid, 490 +/- 50 x 230 +/- 20 µm, membranous, pale brown, semi-transparent when young, darker to almost black when old. Neck not-clearly differentiated, sometime slightly curved towards the light, decorated by small triangular short swollen hairs typical of *Schizothecium* perithecia. Peridium with *textura angularis*. Asci four-spored and claviform. Spores obliquely uniseriate, transversely septate with hyaline primary appendage and secondary appendages at both poles often difficult to see. Spore head 27.0 +/- 1.6 x 14.8 +/- 0.6 µm, ellipsoidal, smooth, flattened at the base, with an apical germ pore. Primary appendage (pedicel) cylindric, slightly tapering towards the apex, 10.5 +/- 1.0 x 3.0 +/- 0.7 µm. Secondary appendage present but evanescent and lacking on most expelled ascospores.

Habitat & Distribution: Saprobic on dung and also likely present in soil, frequently isolated from rabbit dung. The type is from the Netherlands on rabbit dung, the other strains are from France (Northern and Southern parts).

Mating strategy: pseudo-homothallic.

Additional strains : France: PSN350, PSN351, PSN352, PSN535, PSN559, PSN571, PSN667, PSN682, PSN705, PSN706, PSN711, PSN774, PSN777, PSN829, PSN935, PSN1002, PSN1007, PSN1040, PSN1041, PSN1069, PSN1070, PSN1105, PSN1216, PSN1236, PSN1433. Netherlands: CBS255.69.

***Schizothecium tetratetrasporum*** Silar ***sp. nov.*** Fig. 9

**Fig. 9.**
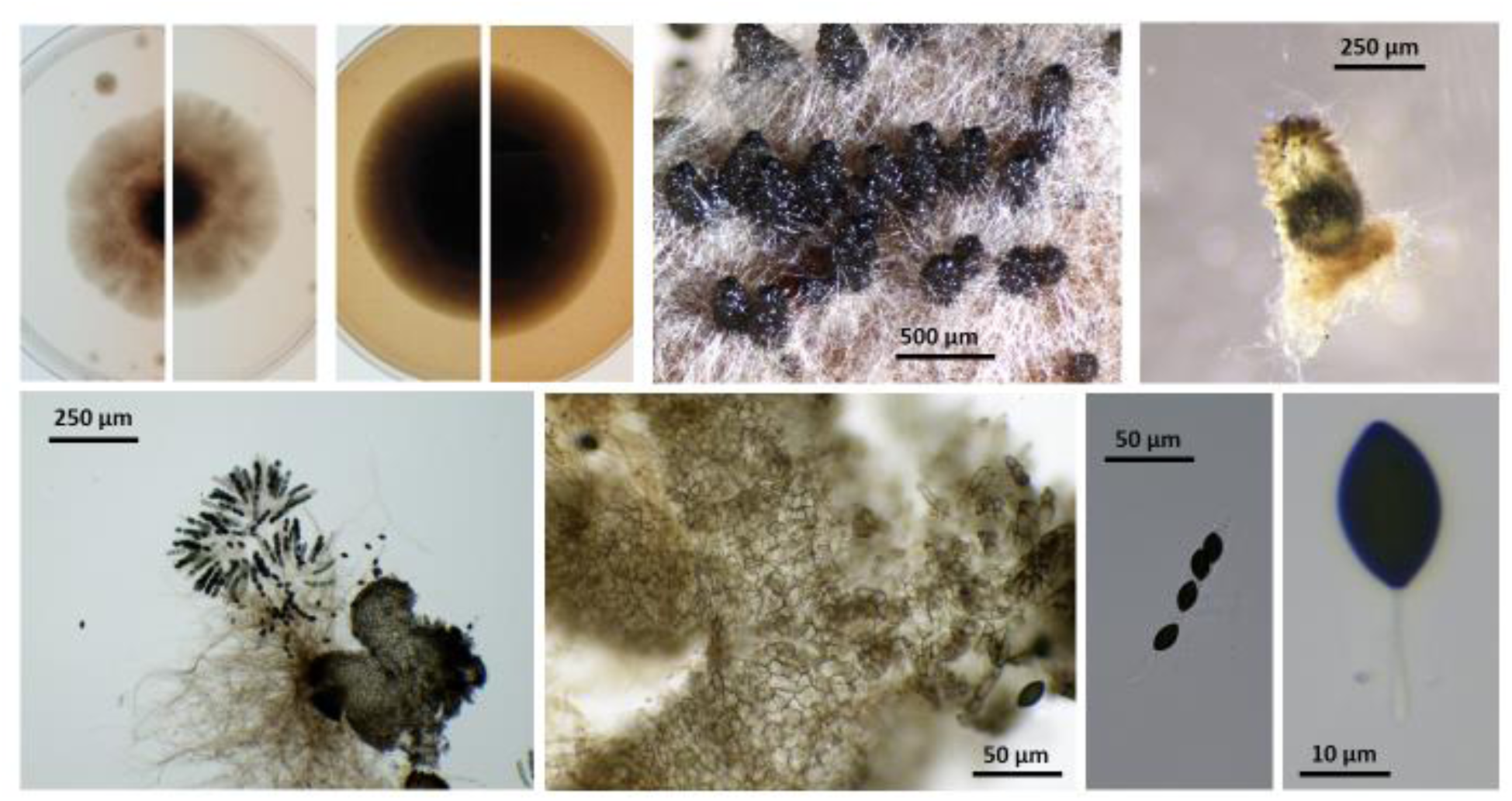
PSN831, the ex-type of *Schizothecium tetratetrasporum*. Top from left to right: culture on M2 top view, culture on M2 reverse view, culture on V8 top view, culture on V8 reverse view, perithecia on M0+ miscanthus, isolated perithecia. Bottom from left to right; squeezed perithecium with rosette of asci and some peridium, enlarge peridium view, ascus and ascospore.

Index Fungorum number:

*Holotype*: PC0798997, ex-type: PSN831

Description: mycelium slow growing (1.0 +/- 0.1 mm/day on M2, 1.4 +/- 0.0 mm day on V8), dark green to almost black with few aerial hyphae on M2 and profuse ones on V8 and often but not always undergoing lysis. Perithecia conical to ovoid, 360 +/- 40 x 220 +/- 20 µm, membranous, pale brown, semi-transparent when young, darker to almost black when old. Neck not-clearly differentiated, sometime slightly curved towards the light, decorated by small triangular short swollen hairs typical of *Schizothecium* perithecia. Peridium with *textura angularis*. Asci four-spored and claviform. Spores obliquely uniseriate, transversely septate with hyaline primary appendage and secondary appendages at both poles often difficult to see. Spore head 24.5 +/- 1.4 x 13.3 +/-0.7 µm, ellipsoidal, smooth, flattened at the base, with an apical germ pore. Primary appendage (pedicel) cylindric, slightly tapering towards the apex, 14.5 +/- 1.4 x 3.0 +/- 0.6 µm. Secondary appendage present but evanescent and lacking on most expelled ascospores.

Habitat & Distribution: Saprobic on dung. Isolated from France (Northern or Southern parts). Mating strategy: pseudo-homothallic.

Additional strains : France: PSN572, PSN708, PSN1067.

***Schizothecium pentatetrasporum*** Silar ***sp. nov.*** Fig. 10

**Fig. 10.**
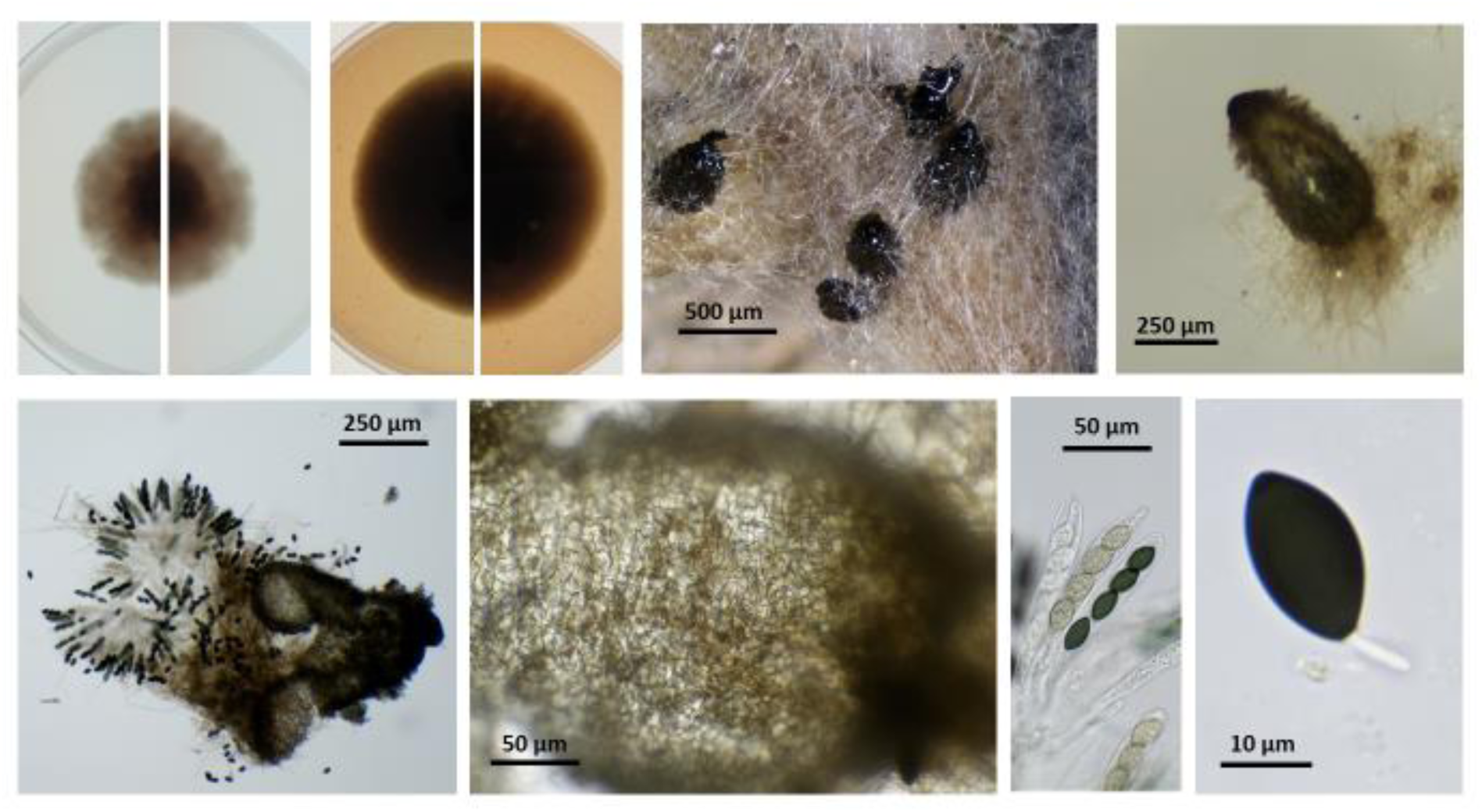
PSN580, the ex-type of *Schizothecium pentatetrasporum*. Top from left to right: culture on M2 top view, culture on M2 reverse view, culture on V8 top view, culture on V8 reverse view, perithecia on M0+ miscanthus, isolated perithecia. Bottom from left to right; squeezed perithecium with rosette of asci and some peridium, enlarge peridium view, asci and ascospore.

Index Fungorum number:

*Holotype*: PC0798999, ex-type: PSN580

Description: mycelium slow growing (1.0 +/- 0.1 mm/day on M2, 1.4 +/- 0.0 mm day on V8), dark green to almost black with few aerial hyphae on M2 and profuse ones on V8 and often but not always undergoing lysis. Perithecia conical to ovoid, 450 +/- 50 x 280 +/- 30 µm, membranous, pale brown, semi-transparent when young, darker to almost black when old. Neck not-clearly differentiated, sometime slightly curved towards the light, decorated by small triangular short swollen hairs typical of *Schizothecium* perithecia. Peridium with *textura angularis*. Asci four-spored and claviform. Spores obliquely uniseriate, transversely septate with hyaline primary appendage and secondary appendages at both poles often difficult to see. Spore head 26.3 +/- 1.2 x 14.2 +/- 0.7 µm, ellipsoidal, smooth, flattened at the base, with an apical germ pore. Primary appendage (pedicel) cylindric, slightly tapering towards the apex, 11.5 +/- 0.6 x 2.7 +/- 0.5 µm. Secondary appendage present but evanescent and lacking on most expelled ascospores.

Habitat & Distribution: Saprobic on dung. Isolated from France (Northern or Southern parts). Mating strategy: pseudo-homothallic.

Additional strains : France: PSN776.

***Schizothecium hexatetrasporum*** Silar ***sp. nov.*** Fig. 11

**Fig. 11.**
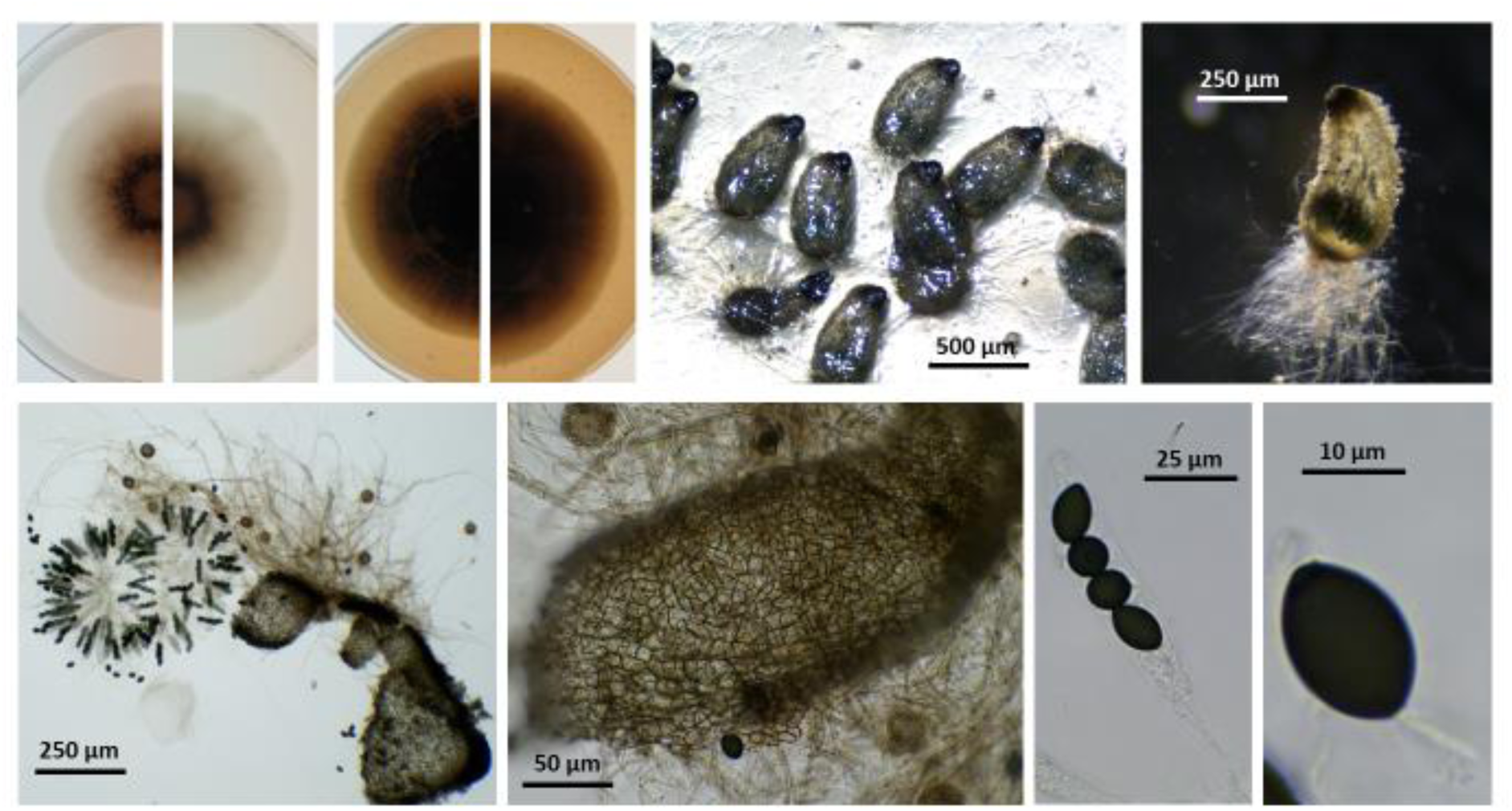
PSN644, the ex-type of *Schizothecium hexatetrasporum*. Top from left to right: culture on M2 top view, culture on M2 reverse view, culture on V8 top view, culture on V8 reverse view, perithecia on M0+ miscanthus, isolated perithecia. Bottom from left to right; squeezed perithecium with rosette of asci and some peridium, enlarge peridium view, ascus and ascospore.

Index Fungorum number:

*Holotype*: PC0799000, ex-type: PSN644

Description: mycelium slow growing (1.1 +/- 0.1 mm/day on M2, 1.4 +/- 0.1 mm day on V8), dark green to almost black with few aerial hyphae on M2 and profuse ones on V8 and often but not always undergoing lysis. Perithecia conical to ovoid, 440 +/- 40 x 230 +/- 20 µm, membranous, pale brown, semi-transparent when young, darker to almost black when old. Neck not-clearly differentiated, sometime slightly curved towards the light, decorated by small triangular short swollen hairs typical of *Schizothecium* perithecia. Peridium with *textura angularis*. Asci four-spored and claviform. Spores obliquely uniseriate, transversely septate with hyaline primary appendage and secondary appendages at both poles often difficult to see. Spore head 24.1 +/- 1.2 x 14.5 +/- 0.9 µm, ellipsoidal, smooth, flattened at the base, with an apical germ pore. Primary appendage (pedicel) cylindric, slightly tapering towards the apex, 10.9 +/- 1.1 x 2.7 +/- 0.4 µm. Secondary appendage present but evanescent and lacking on most expelled ascospores.

Habitat & Distribution: Saprobic on dung. Isolated from Ontario, Canada. Only one strain was isolated for this species from sylvilagus dung collected in Canatara park in Ontario.

Mating strategy: pseudo-homothallic.

***Schizothecium heptatetrasporum*** Silar ***sp. nov.*** Fig. 12

**Fig. 12.**
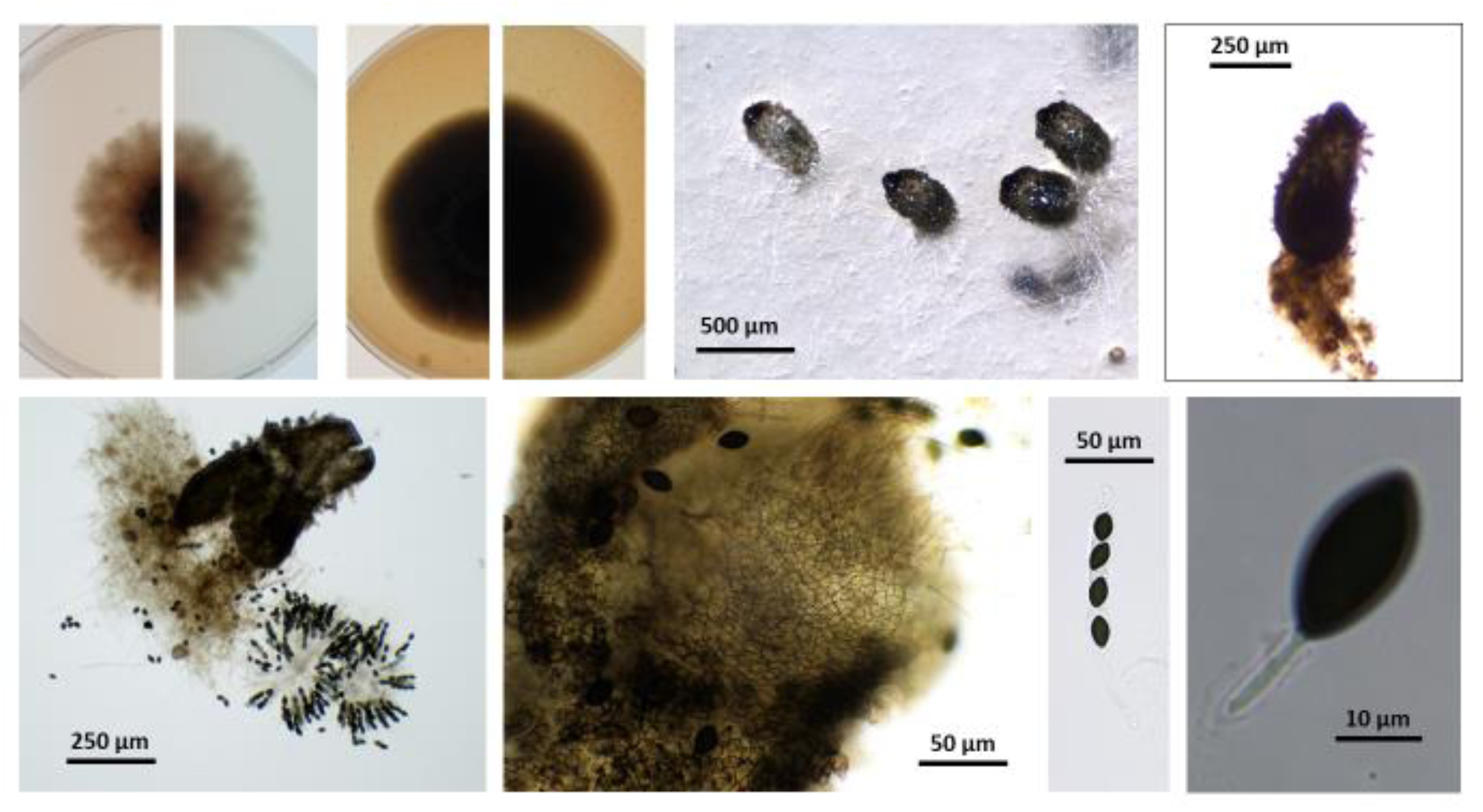
PSN590, the ex-type of *Schizothecium heptatetrasporum*. Top from left to right: culture on M2 top view, culture on M2 reverse view, culture on V8 top view, culture on V8 reverse view, perithecia on M0+ miscanthus, isolated perithecia. Bottom from left to right; squeezed perithecium with rosette of asci and some peridium, enlarge peridium view, ascus and ascospore.

Index Fungorum number:

*Holotype*: PC0799001, ex-type: PSN590

Description: mycelium slow growing (1.0 +/- 0.1 mm/day on M2, 1.2 +/- 0.0 mm day on V8), dark green to almost black with few aerial hyphae on M2 and profuse ones on V8 and often but not always undergoing lysis. Perithecia conical to ovoid, 460 +/- 40 x 230 +/- 20 µm, membranous, pale brown, semi-transparent when young, darker to almost black when old. Neck not-clearly differentiated, sometime slightly curved towards the light, decorated by small triangular short swollen hairs typical of *Schizothecium* perithecia. Peridium with *textura angularis*. Asci four-spored and claviform. Spores obliquely uniseriate, transversely septate with hyaline primary appendage and secondary appendages at both poles often difficult to see. Spore head 25.6 +/- 2.0 x 15.3 +/-1.2 µm, ellipsoidal, smooth, flattened at the base, with an apical germ pore. Primary appendage (pedicel) cylindric, slightly tapering towards the apex, 12.0 +/- 1.9 x 2.4 +/- 0.5 µm. Secondary appendage present but evanescent and lacking on most expelled ascospores.

Habitat & Distribution: Saprobic on dung. Isolated from France. Only one strain was isolated for this species from hare dung collected in the “Alpes de Haute provence” from Southern France.

Mating strategy: pseudo-homothallic.

***Schizothecium octatetrasporum*** Silar ***sp. nov.*** Fig. 13

**Fig. 13.**
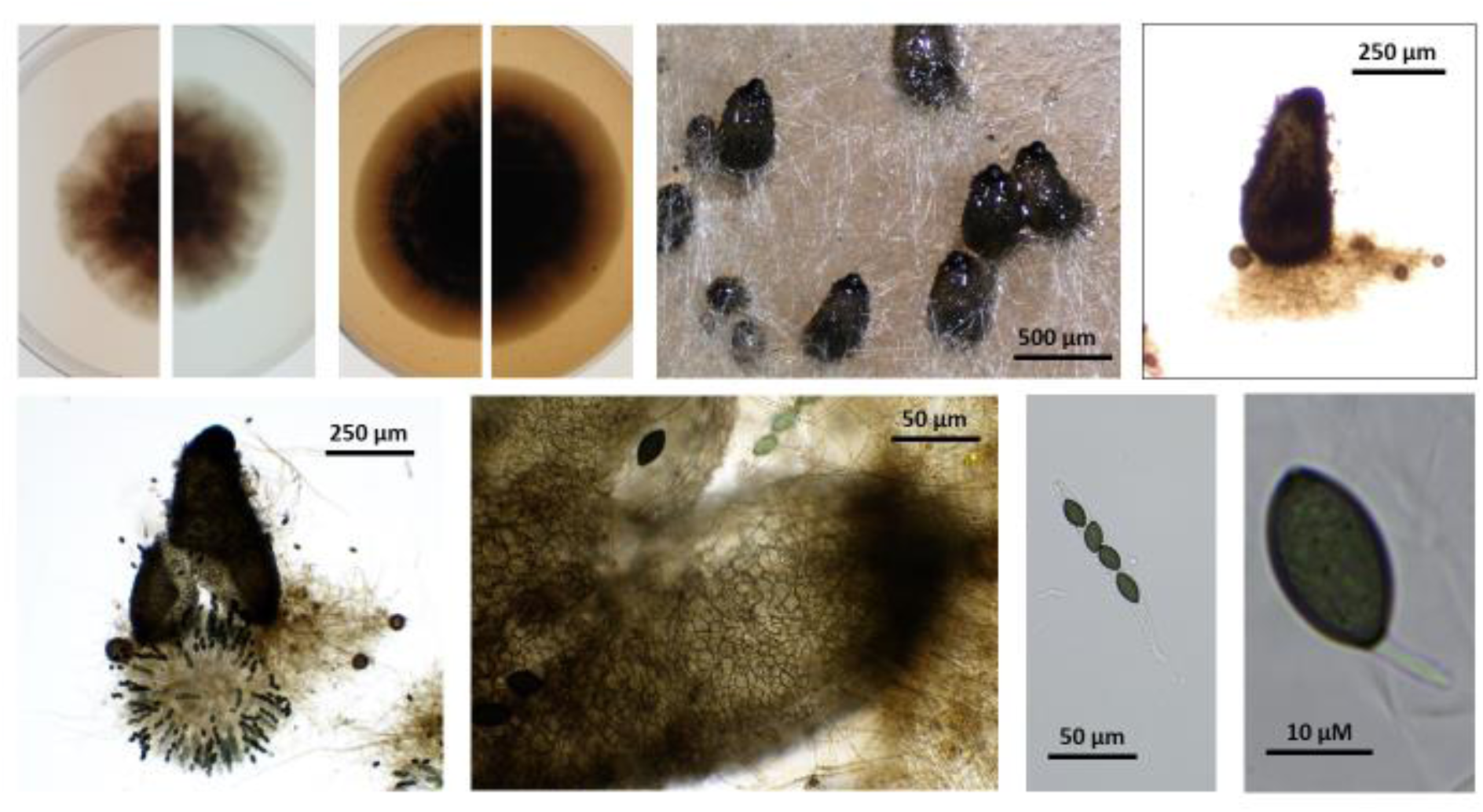
PSN663, the ex-type of *Schizothecium octatetrasporum*. Top from left to right: culture on M2 top view, culture on M2 reverse view, culture on V8 top view, culture on V8 reverse view, perithecia on M0+ miscanthus, isolated perithecia. Bottom from left to right; squeezed perithecium with rosette of asci and some peridium, enlarge peridium view, ascus and ascospore.

Index Fungorum number:

*Holotype*: PC0799002, ex-type: PSN663

Description: mycelium slow growing (1.3 +/- 0.1 mm/day on M2, 1.5 +/- 0.0 mm day on V8), dark green to almost black with few aerial hyphae on M2 and profuse ones on V8 and often but not always undergoing lysis. Perithecia conical to ovoid, 510 +/- 44 x 250 +/- 10 µm, membranous, pale brown, semi-transparent when young, darker to almost black when old. Neck not-clearly differentiated, sometime slightly curved towards the light, decorated by small triangular short swollen hairs typical of *Schizothecium* perithecia. Peridium with *textura angularis*. Asci four-spored and claviform. Spores obliquely uniseriate, transversely septate with hyaline primary appendage and secondary appendages at both poles often difficult to see. Spore head 26.4 +/-1.4 x 15.0 +/- 0.4 µm, ellipsoidal, smooth, flattened at the base, with an apical germ pore. Primary appendage (pedicel) cylindric, slightly tapering towards the apex, 11.1+/- 1.4 x 2.4 +/- 0.3 µm. Secondary appendage present but evanescent and lacking on most expelled ascospores.

Habitat & Distribution: Saprobic on dung. Isolated from Ontario, Canada. Only one strain was isolated for this species from sylvilagus dung collected in Elgin County, Ontario, Canada.

Mating strategy: pseudo-homothallic.

Note: The sole strain available for this species has a 0.8% divergence with the strains of *S. ditetrasporum* making them the two most closely related species (Fig. 4). However, they share very few highly conserved sequences (about 5-6% of sequences with less than 1‰ divergence) showing that the *S. octotetrasporum* and *S. ditetrasporum* do not frequently cross in the wild.

***Schizothecium enneatetrasporum*** Silar ***sp. nov.*** Fig. 14

**Fig. 14.**
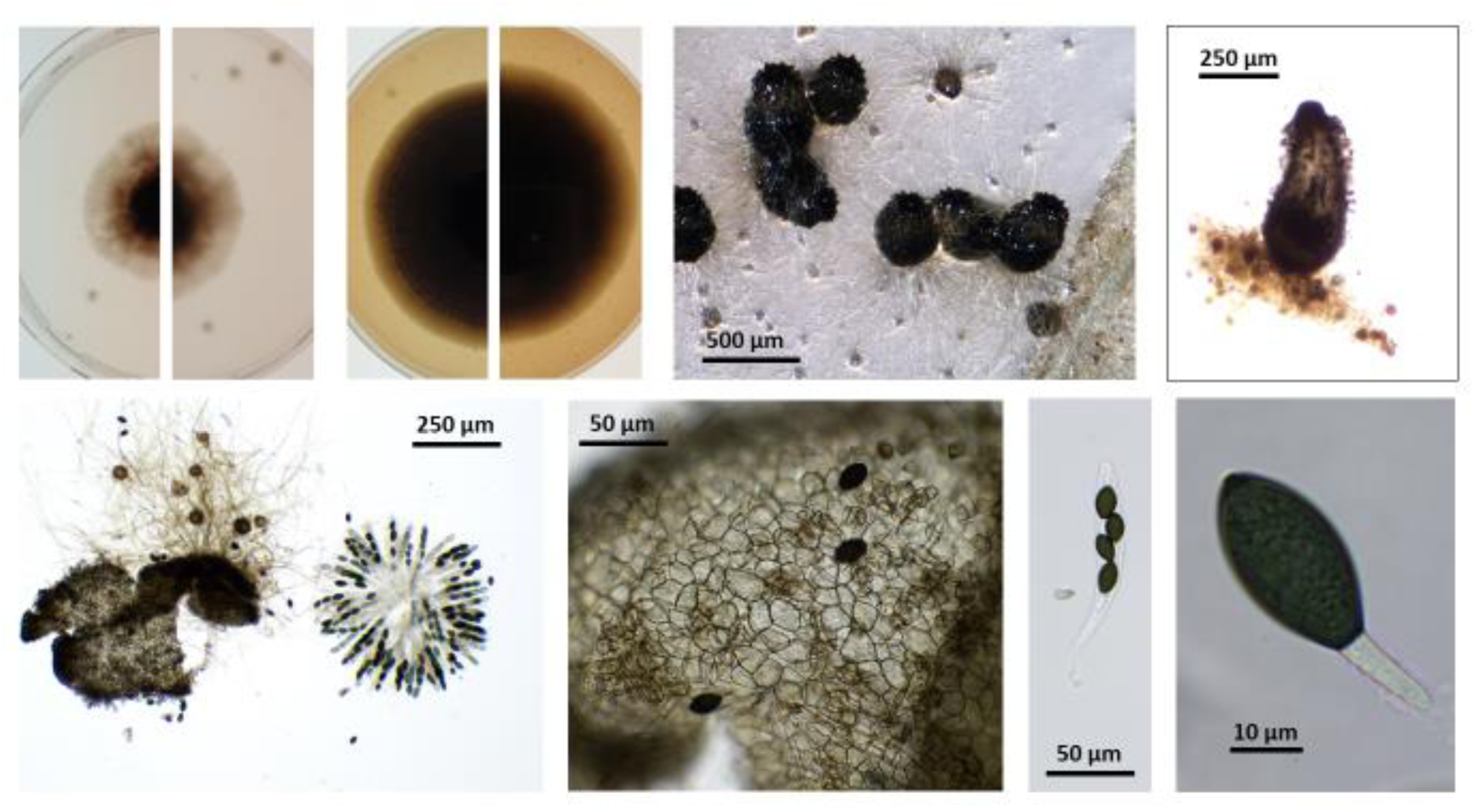
PSN1042, the ex-type of *Schizothecium enneatetrasporum*. Top from left to right: culture on M2 top view, culture on M2 reverse view, culture on V8 top view, culture on V8 reverse view, perithecia on M0+ miscanthus, isolated perithecia. Bottom from left to right; squeezed perithecium with rosette of asci and some peridium, enlarge peridium view, ascus and ascospore.

Index Fungorum number:

*Holotype*: PC0799003, ex-type: PSN1042

Description: mycelium slow growing (1.1 +/- 0.0 mm/day on M2, 1.4 +/- 0.1 mm day on V8), dark green to almost black with few aerial hyphae on M2 and profuse ones on V8 and often but not always undergoing lysis. Perithecia conical to ovoid, 520 +/- 50 x 230 +/- 20 µm, membranous, pale brown, semi-transparent when young, darker to almost black when old. Neck not-clearly differentiated, sometime slightly curved towards the light, decorated by small triangular short swollen hairs typical of *Schizothecium* perithecia. Peridium with *textura angularis*. Asci four-spored and claviform. Spores obliquely uniseriate, transversely septate with hyaline primary appendage and secondary appendages at both poles often difficult to see. Spore head 26.4 +/- 1.0 x 15.2 +/- 0.5 µm, ellipsoidal, smooth, flattened at the base, with an apical germ pore. Primary appendage (pedicel) cylindric, slightly tapering towards the apex, 11.2 +/- 1.8 x 2.7 +/- 0.4 µm. Secondary appendage present but evanescent and lacking on most expelled ascospores.

Habitat & Distribution: Saprobic on dung and possibly in soil. Isolated from France and New Zealand. The type strain for this species was isolated from rabbit dung collected in Brittany, France.

Mating strategy: pseudo-homothallic.

Additional strains : France: PSN561, PSN1219; New Zealand: CBS132.94.

***Schizothecium decatetrasporum*** Silar ***sp. nov.*** Fig. 15

**Fig. 15.**
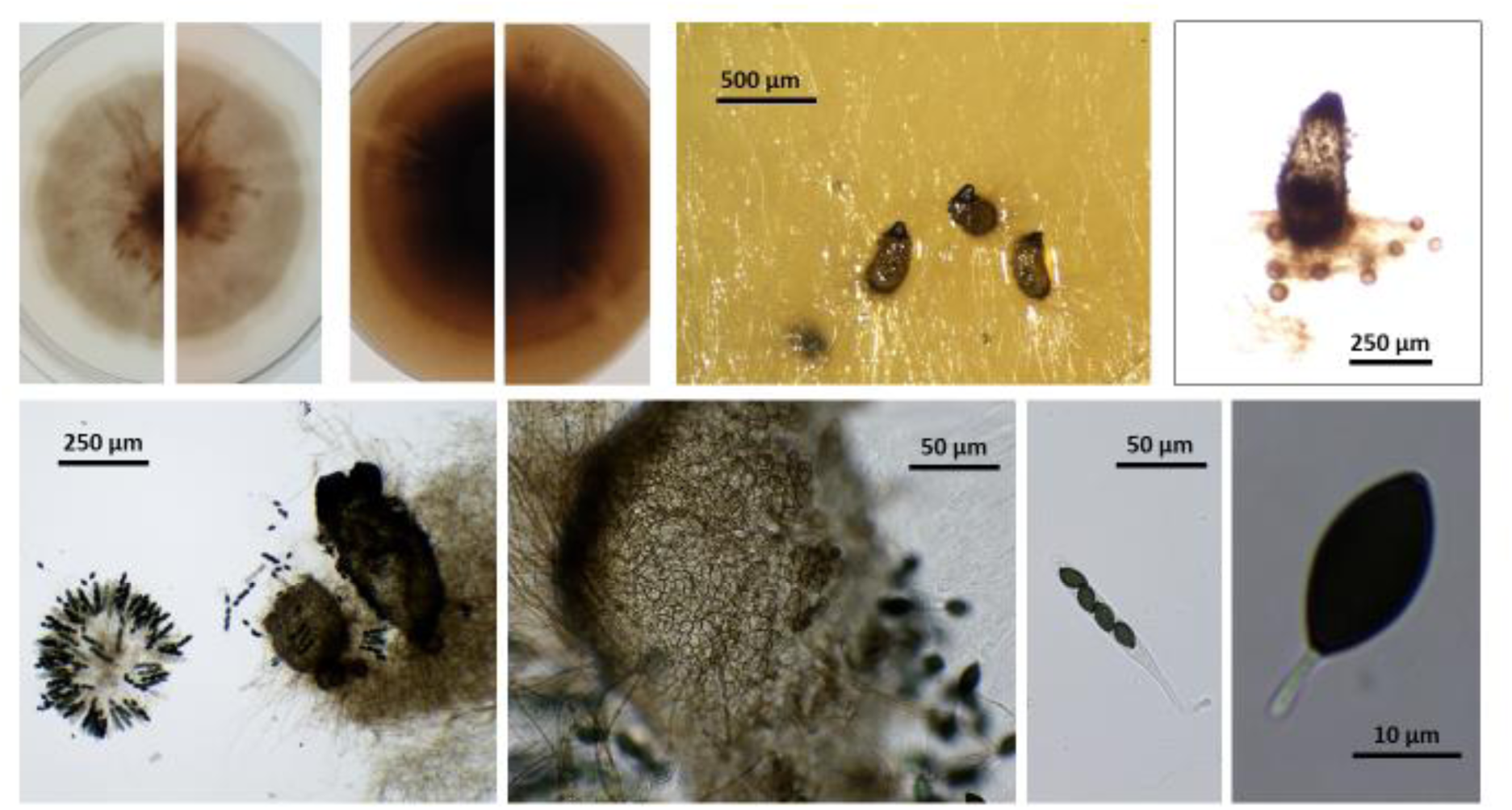
PSN959, the ex-type of *Schizothecium decatetrasporum*. Top from left to right: culture on M2 top view, culture on M2 reverse view, culture on V8 top view, culture on V8 reverse view, perithecia on M0+ miscanthus, isolated perithecia. Bottom from left to right; squeezed perithecium with rosette of asci and some peridium, enlarge peridium view, ascus and ascospore.

Index Fungorum number:

*Holotype*: PC0799004, ex-type: PSN959

Description: mycelium slow growing (1.1 +/- 0.0 mm/day on M2, 1.1 +/- 0.1 mm day on V8), dark green to almost black with few aerial hyphae on M2 and profuse ones on V8 and often but not always undergoing lysis. Perithecia conical to ovoid, 430 +/- 30 x 210 +/- 10 µm, membranous, pale brown, semi-transparent when young, darker to almost black when old. Neck not-clearly differentiated, sometime slightly curved towards the light, decorated by small triangular short swollen hairs typical of *Schizothecium* perithecia. Peridium with *textura angularis*. Asci four-spored and claviform. Spores obliquely uniseriate, transversely septate with hyaline primary appendage and secondary appendages at both poles often difficult to see. Spore head 26.3 +/- 1.1 x 13.6 +/-0.7 µm, ellipsoidal, smooth, flattened at the base, with an apical germ pore. Primary appendage (pedicel) cylindric, slightly tapering towards the apex, 12.5 +/- 1.7 x 2.6 +/- 0.5 µm. Secondary appendage present but evanescent and lacking on most expelled ascospores.

Habitat & Distribution: Saprobic on dung. Isolated from France and Italy in the Alps mountain. The type strain for this species was isolated from boar dung collected in Vercors regional park in the Alps in Southern France.

Mating strategy: pseudo-homothallic. Additional strains : Italy: PSN302.

***Schizothecium hendecatetrasporum*** Silar ***sp. nov.*** Fig. 16

**Fig. 16.**
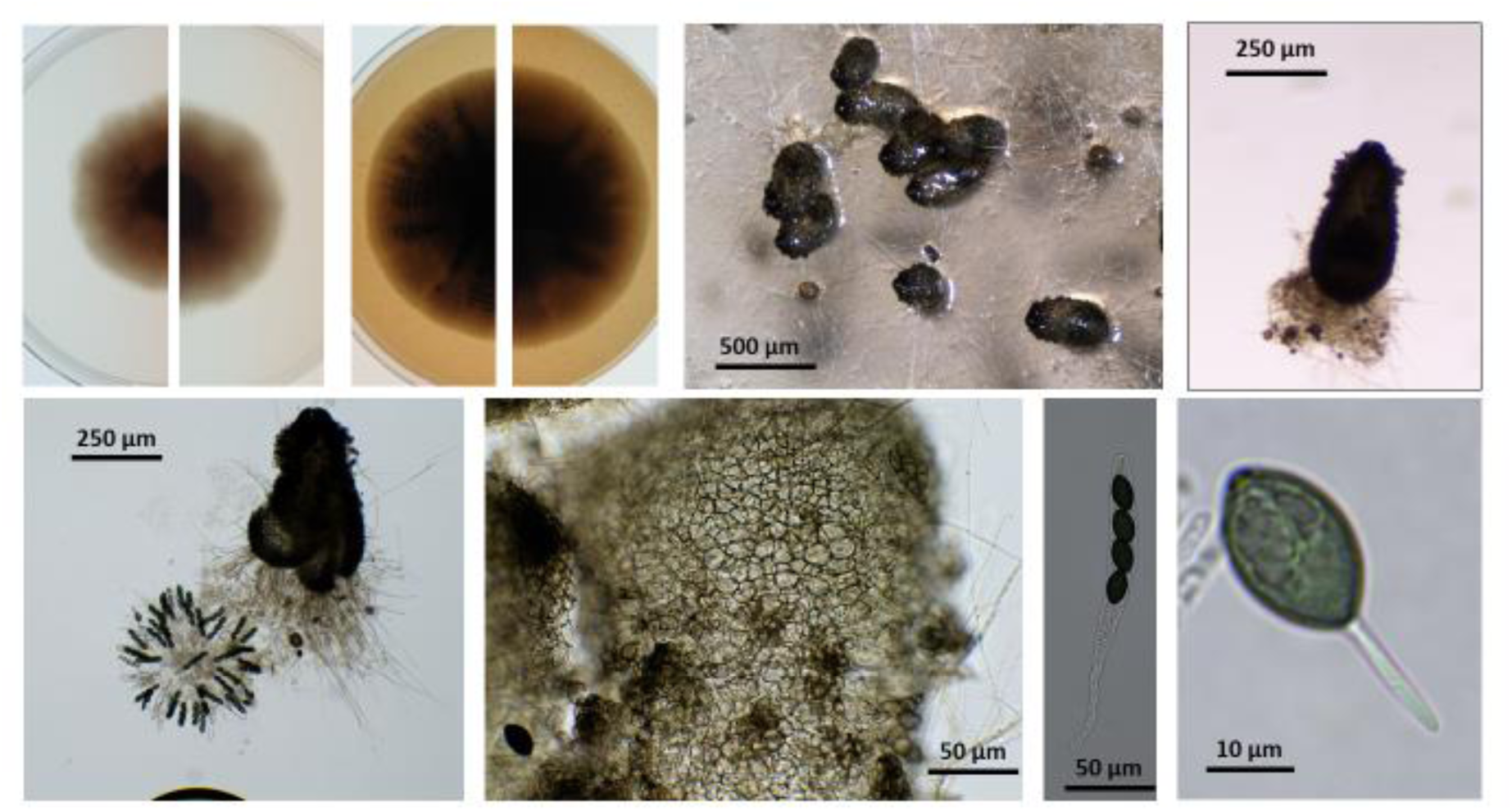
PSN1310, the ex-type of *Schizothecium hendecatetrasporum*. Top from left to right: culture on M2 top view, culture on M2 reverse view, culture on V8 top view, culture on V8 reverse view, perithecia on M0+ miscanthus, isolated perithecia. Bottom from left to right; squeezed perithecium with rosette of asci and some peridium, enlarge peridium view, ascus and ascospore.

Index Fungorum number:

*Holotype*: PC0799005, ex-type: PSN310

Description: mycelium slow growing (1.0 +/- 0.1 mm/day on M2, 1.4 +/- 0.1 mm day on V8), dark green to almost black with few aerial hyphae on M2 and profuse ones on V8 and often but not always undergoing lysis. Perithecia conical to ovoid, 450 +/- 41 x 210 +/- 10 µm, membranous, pale brown, semi-transparent when young, darker to almost black when old. Neck not-clearly differentiated, sometime slightly curved towards the light, decorated by small triangular short swollen hairs typical of *Schizothecium* perithecia. Peridium with *textura angularis*. Asci four-spored and claviform. Spores obliquely uniseriate, transversely septate with hyaline primary appendage and secondary appendages at both poles often difficult to see. Spore head 26.9 +/- 0.9 x 15.3 +/- 0.6 µm, ellipsoidal, smooth, flattened at the base, with an apical germ pore. Primary appendage (pedicel) cylindric, slightly tapering towards the apex, 13.0 +/- 1.5 x 3.2 +/- 0.6 µm. Secondary appendage present but evanescent and lacking on most expelled ascospores.

Habitat & Distribution: Saprobic on dung. Isolated from central Chile. Only one strain for this species was isolated from rabbit dung collected near Santiago Chile.

Mating strategy: pseudo-homothallic.

***Schizothecium pseudotetrasporum*** Silar ***sp. nov.*** Fig. 17

**Fig. 17.**
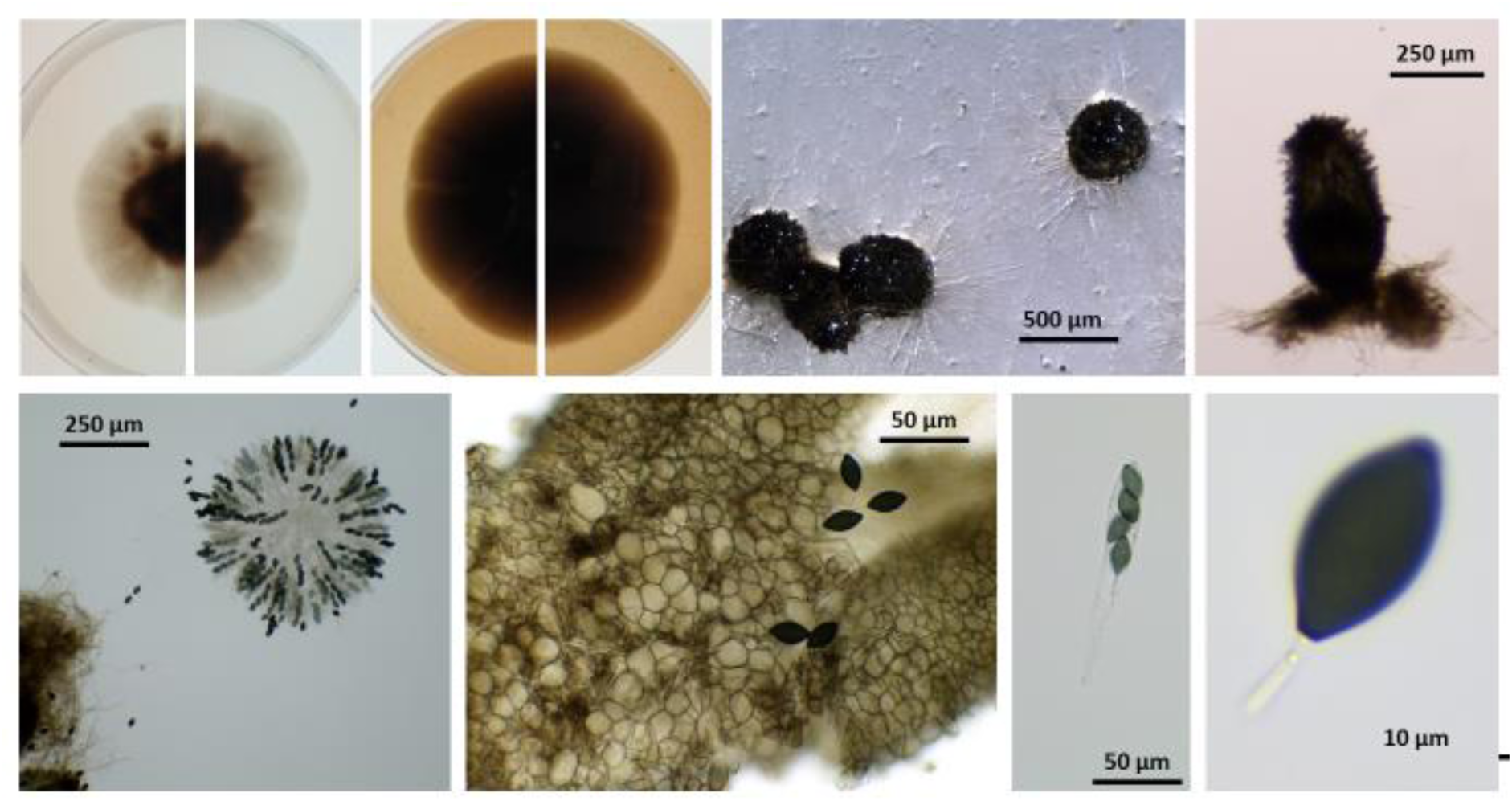
PSN936, the ex-type of *Schizothecium pseudotetrasporum*. Top from left to right: culture on M2 top view, culture on M2 reverse view, culture on V8 top view, culture on V8 reverse view, perithecia on M0+ miscanthus, isolated perithecia. Bottom from left to right; squeezed perithecium with rosette of asci and some peridium, enlarge peridium view, ascus and ascospore.

Index Fungorum number:

*Holotype*: PC0799006, ex-type: PSN936

Description: mycelium slow growing (1.1 +/- 0.1 mm/day on M2, 1.4 +/- 0.1 mm day on V8), dark green to almost black with few aerial hyphae on M2 and profuse ones on V8 and often but not always undergoing lysis. Perithecia conical to ovoid, 490 +/- 40 x 260 +/- 30 µm, membranous, pale brown, semi-transparent when young, darker to almost black when old. Neck not-clearly differentiated, sometime slightly curved towards the light, decorated by small triangular short swollen hairs typical of *Schizothecium* perithecia. Peridium with *textura angularis*. Asci four-spored and claviform. Spores obliquely uniseriate, transversely septate with hyaline primary appendage and secondary appendages at both poles often difficult to see. Spore head 26.0 +/- 1.1 x 14.4 +/- 0.5 µm, ellipsoidal, smooth, flattened at the base, with an apical germ pore. Primary appendage (pedicel) cylindric, slightly tapering towards the apex, 8.8 +/- 1.4 x 2.6 +/- 0.2 µm. Secondary appendage present but evanescent and lacking on most expelled ascospores.

Habitat & Distribution: Saprobic in soil as the two strains newly obtained for this species were both from soil and the one available in collection (IMI320477) was found growing on *Tuber rufum* fruit body. Isolated from France and the U.K. The type strain for this species was isolated from a soil collected in the Alps in Southern France.

Mating strategy: pseudo-homothallic.

Additional strains : France: PSN518. U.K. : IMI320477.

***Schizothecium dipseudotetrasporum*** Silar ***sp. nov.*** Fig. 18

**Fig. 18.**
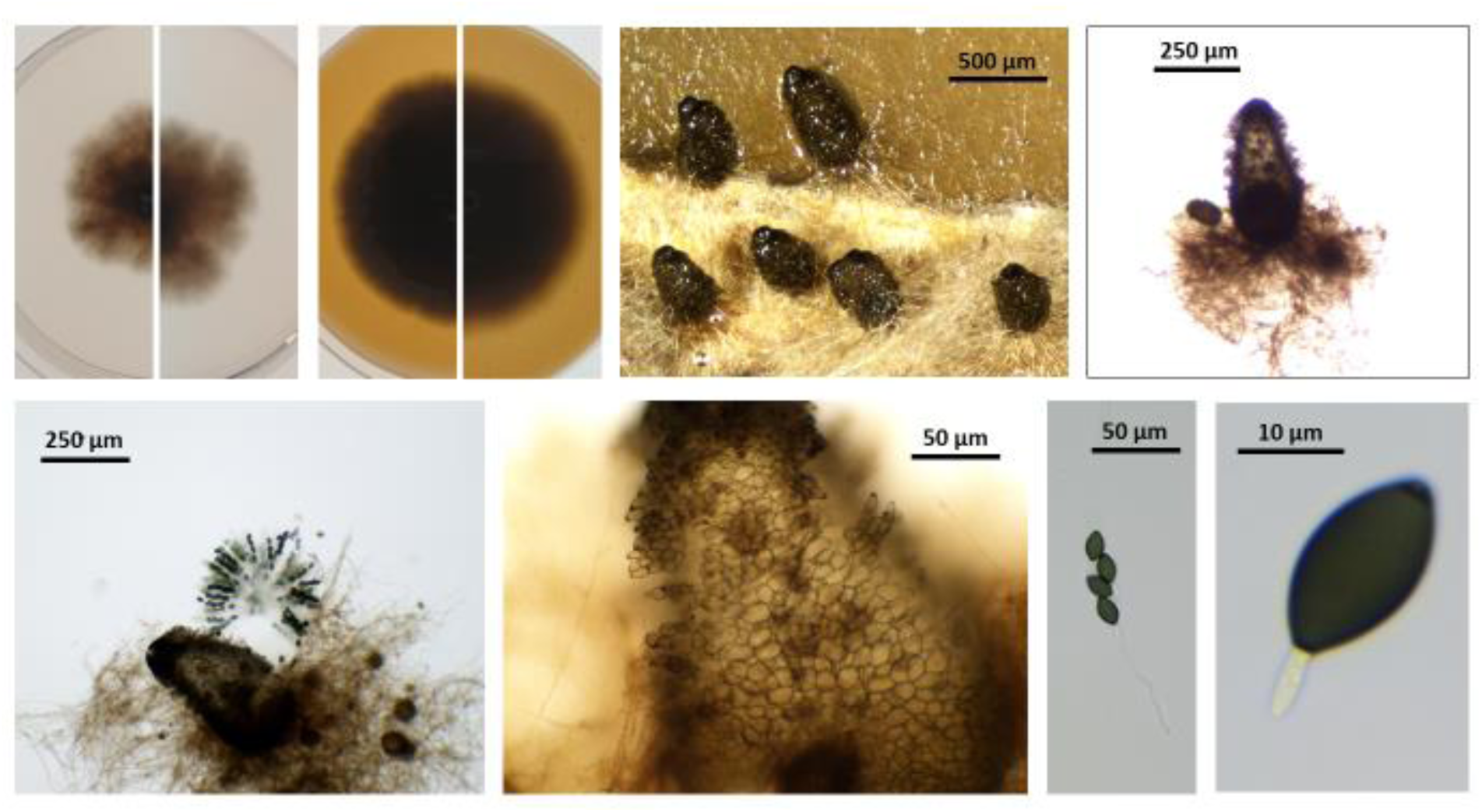
PSQ32, the ex-type of *Schizothecium dipseudotetrasporum*. Top from left to right: culture on M2 top view, culture on M2 reverse view, culture on V8 top view, culture on V8 reverse view, perithecia on M0+ miscanthus, isolated perithecia. Bottom from left to right; squeezed perithecium with rosette of asci and some peridium, enlarge peridium view, ascus and ascospore.

Index Fungorum number:

*Holotype*: PC0799007, ex-type: PSQ32

Description: mycelium slow growing (1.1 +/- 0.1 mm/day on M2, 1.4 +/- 0.1 mm day on V8), dark green to almost black with few aerial hyphae on M2 and profuse ones on V8 and often but not always undergoing lysis. Perithecia conical to ovoid, 370 +/- 40 x 200 +/- 10 µm, membranous, pale brown, semi-transparent when young, darker to almost black when old. Neck not-clearly differentiated, sometime slightly curved towards the light, decorated by small triangular short swollen hairs typical of *Schizothecium* perithecia. Peridium with *textura angularis*. Asci four-spored and claviform. Spores obliquely uniseriate, transversely septate with hyaline primary appendage and secondary appendages at both poles often difficult to see. Spore head 24.4 +/- 1.5 x 13.9 +/- 0.5, ellipsoidal, smooth, flattened at the base, with an apical germ pore. Primary appendage (pedicel) cylindric, slightly tapering towards the apex, 10.5 +/- 1.4 x 2.9 +/- 0.3 µm. Secondary appendage present but evanescent and lacking on most expelled ascospores.

Habitat & Distribution: Saprobic on dung. Isolated from Quebec, Canada. The only available strain for this species was isolated from a moose dung collected in the Anticosti park in Quebec, Canada.

Mating strategy: pseudo-homothallic.

***Schizothecium tripseudotetrasporum*** Silar ***sp. nov.*** Fig. 19

**Fig. 19.**
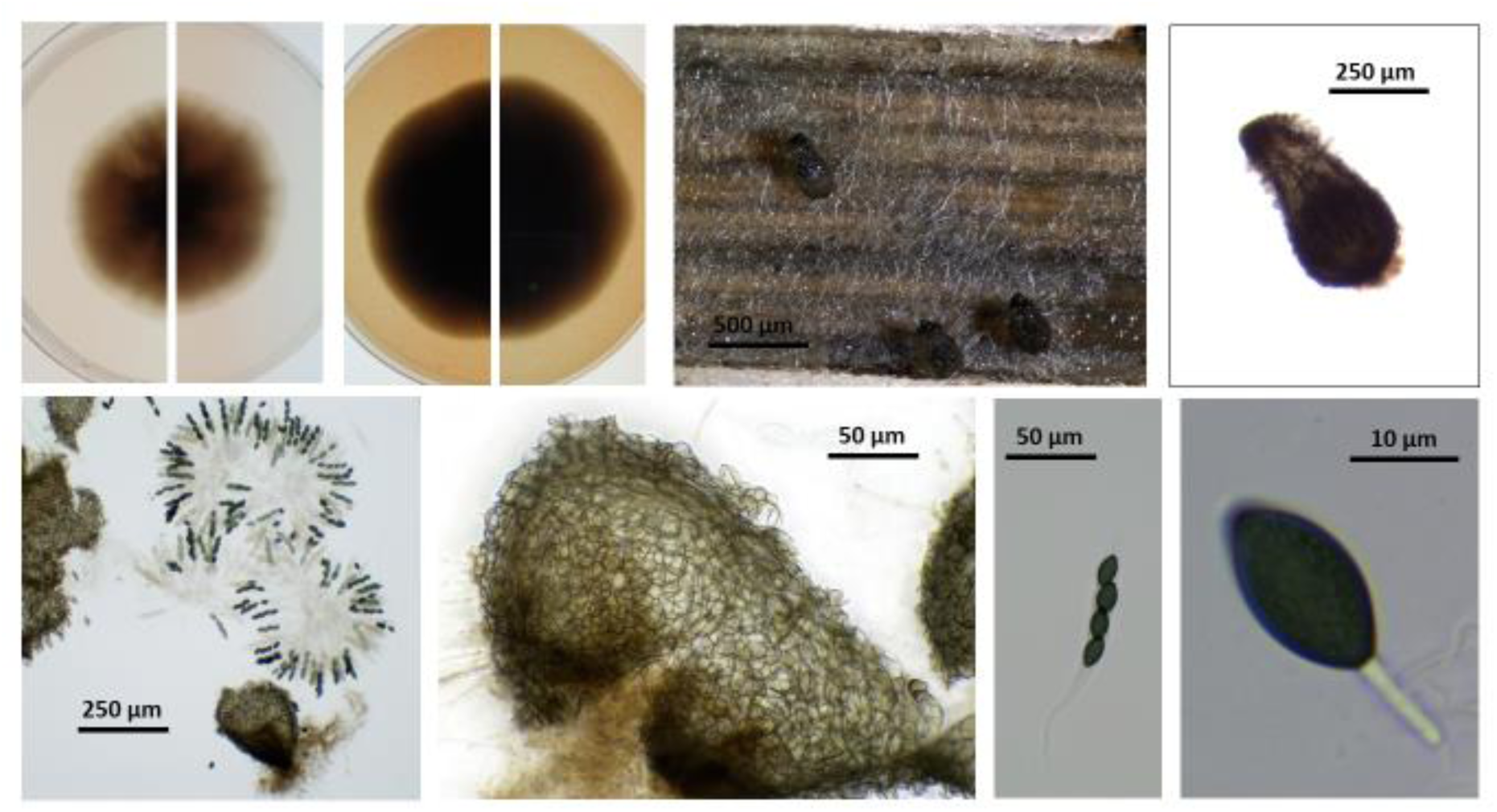
PSN707, the ex-type of *Schizothecium tripseudotetrasporum*. Top from left to right: culture on M2 top view, culture on M2 reverse view, culture on V8 top view, culture on V8 reverse view, perithecia on M0+ miscanthus, isolated perithecia. Bottom from left to right; squeezed perithecium with rosette of asci and some peridium, enlarge peridium view, ascus and ascospore.

Index Fungorum number:

*Holotype*: PC0799008, ex-type: PSN707

Description: mycelium slow growing (1.3 +/- 0.1 mm/day on M2, 1.4 +/- 0.1 mm day on V8), dark green to almost black with few aerial hyphae on M2 and profuse ones on V8 and often but not always undergoing lysis. Perithecia conical to ovoid, 450 +/- 40 x 234 +/- 30 µm, membranous, pale brown, semi-transparent when young, darker to almost black when old. Neck not-clearly differentiated, sometime slightly curved towards the light, decorated by small triangular short swollen hairs typical of *Schizothecium* perithecia. Peridium with *textura angularis*. Asci four-spored and claviform. Spores obliquely uniseriate, transversely septate with hyaline primary appendage and secondary appendages at both poles often difficult to see. Spore head 25.0 +/- 1.0 x 13.8 +/- 0.6, ellipsoidal, smooth, flattened at the base, with an apical germ pore. Primary appendage (pedicel) cylindric, slightly tapering towards the apex, 7.8 +/- 1.1 x 2.4 +/- 0.5 µm. Secondary appendage present but evanescent and lacking on most expelled ascospores.

Habitat & Distribution: Saprobic on dung. Isolated from Ontario, Canada. The only available strain for this species was isolated from a sylvilagus dung collected near Toronto, Ontario, Canada.

Mating strategy: pseudo-homothallic.

***Schizothecium tetrapseudotetrasporum*** Silar ***sp. nov.*** Fig. 20

**Fig. 20.**
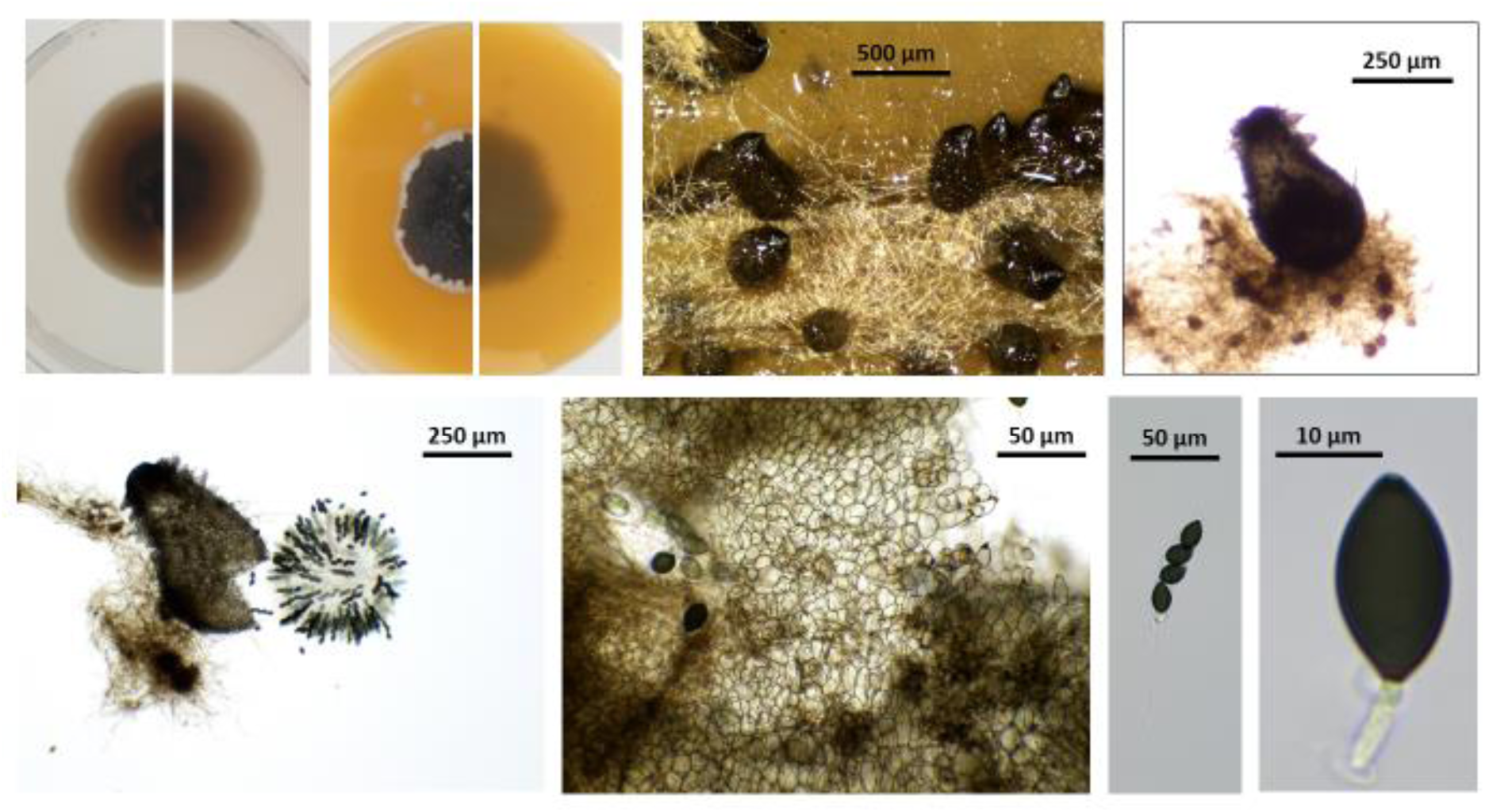
PSN684, the ex-type of *Schizothecium tetrapseudotetrasporum*. Top from left to right: culture on M2 top view, culture on M2 reverse view, culture on V8 top view, culture on V8 reverse view, perithecia on M0+ miscanthus, isolated perithecia. Bottom from left to right; squeezed perithecium with rosette of asci and some peridium, enlarge peridium view, ascus and ascospore.

Index Fungorum number:

*Holotype*: PC0799009, ex-type: PSN684

Description: mycelium slow growing (1.1 +/- 0.0 mm/day on M2, 1.4 +/- 0.1 mm day on V8), dark green to almost black with few aerial hyphae on M2 and profuse ones on V8 and often but not always undergoing lysis. Perithecia conical to ovoid, 400 +/- 50 x 250 +/- 30 µm, membranous, pale brown, semi-transparent when young, darker to almost black when old. Neck not-clearly differentiated, sometime slightly curved towards the light, decorated by small triangular short swollen hairs typical of *Schizothecium* perithecia. Peridium with *textura angularis*. Asci four-spored and claviform. Spores obliquely uniseriate, transversely septate with hyaline primary appendage and secondary appendages at both poles often difficult to see. Spore head 26.4 +/- 1.4 x 14.2 +/- 0.6, ellipsoidal, smooth, flattened at the base, with an apical germ pore. Primary appendage (pedicel) cylindric, slightly tapering towards the apex, 13.8 +/- 1.3 x 2.7 +/- 0.3 µm. Secondary appendage present but evanescent and lacking on most expelled ascospores.

Habitat & Distribution: Saprobic on dung and possibly in soil. Isolated from France (Northern and Southern part) and Canada.

Mating strategy: pseudo-homothallic.

Additional strains : Canada: PSN709. France : PSN584, PSN668.

***Schizothecium pentapseudotetrasporum*** Silar ***sp. nov.*** Fig. 21

**Fig. 21.**
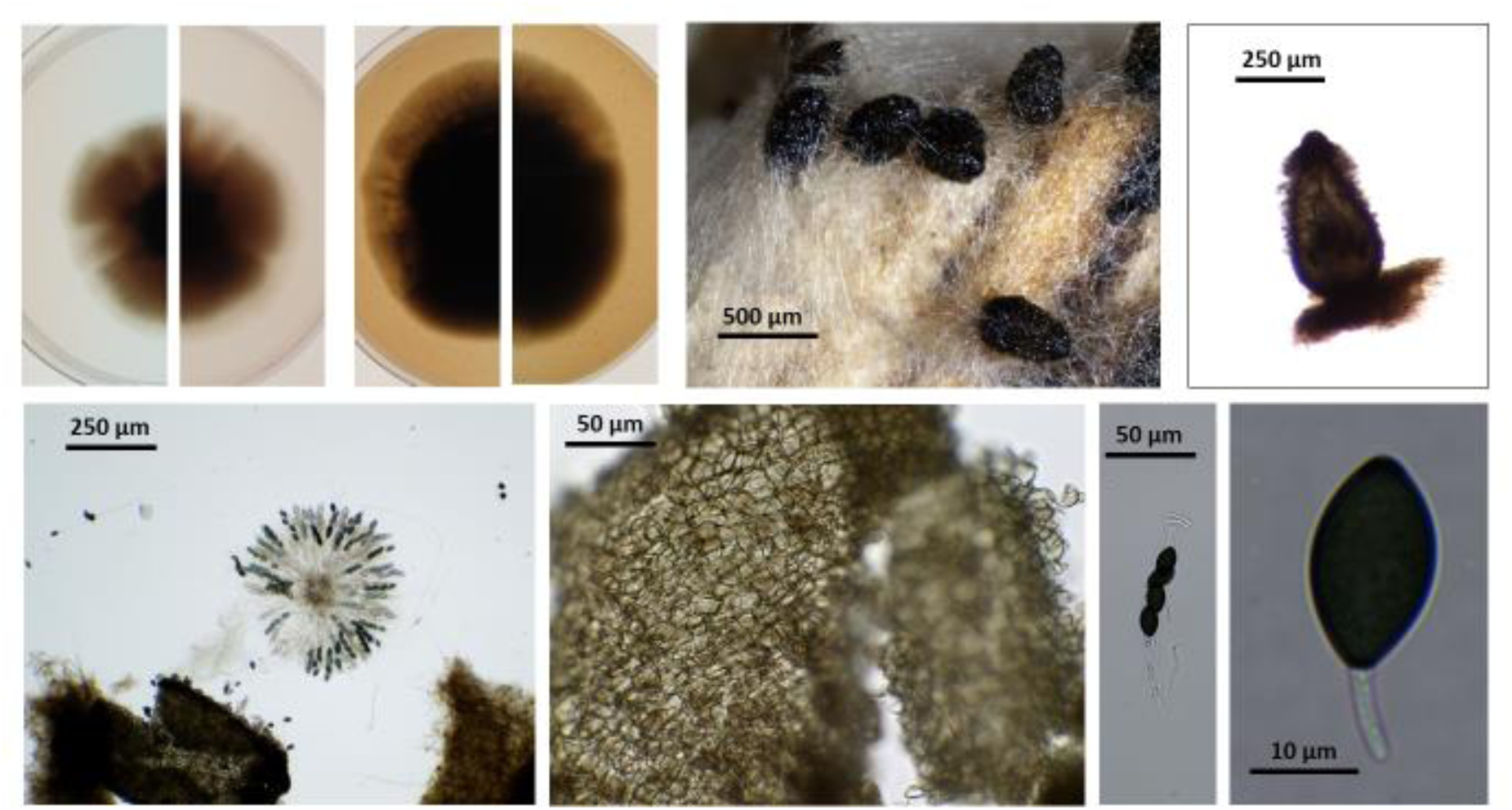
PSN377, the ex-type of *Schizothecium pentapseudotetrasporum*. Top from left to right: culture on M2 top view, culture on M2 reverse view, culture on V8 top view, culture on V8 reverse view, perithecia on M0+ miscanthus, isolated perithecia. Bottom from left to right; squeezed perithecium with rosette of asci and some peridium, enlarge peridium view, ascus and ascospore.

Index Fungorum number:

*Holotype*: PC0799010, ex-type: PSN377

Description: mycelium slow growing (1.2 +/- 0.1 mm/day on M2, 1.6 +/- 0.1 mm day on V8), dark green to almost black with few aerial hyphae on M2 and profuse ones on V8 and often but not always undergoing lysis. Perithecia conical to ovoid, 480 +/- 40 x 270 +/- 20 µm, membranous, pale brown, semi-transparent when young, darker to almost black when old. Neck not-clearly differentiated, sometime slightly curved towards the light, decorated by small triangular short swollen hairs typical of *Schizothecium* perithecia. Peridium with *textura angularis*. Asci four-spored and claviform. Spores obliquely uniseriate, transversely septate with hyaline primary appendage and secondary appendages at both poles often difficult to see. Spore head 26.9 +/- 2.1 x 14.8 +/- 1.1, ellipsoidal, smooth, flattened at the base, with an apical germ pore. Primary appendage (pedicel) cylindric, slightly tapering towards the apex, 10.2 +/- 1.5 x 2.3 +/- 0.4 µm. Secondary appendage present but evanescent and lacking on most expelled ascospores.

Habitat & Distribution: Saprobic on dung. Isolated from Quebec, Canada. The only strain available for this species was isolated from moose dung collected in Anticosti, Quebec, Canada.

Mating strategy: pseudo-homothallic.

***Schizothecium octosporum*** Silar ***sp. nov.*** Fig. 22

**Fig. 22.**
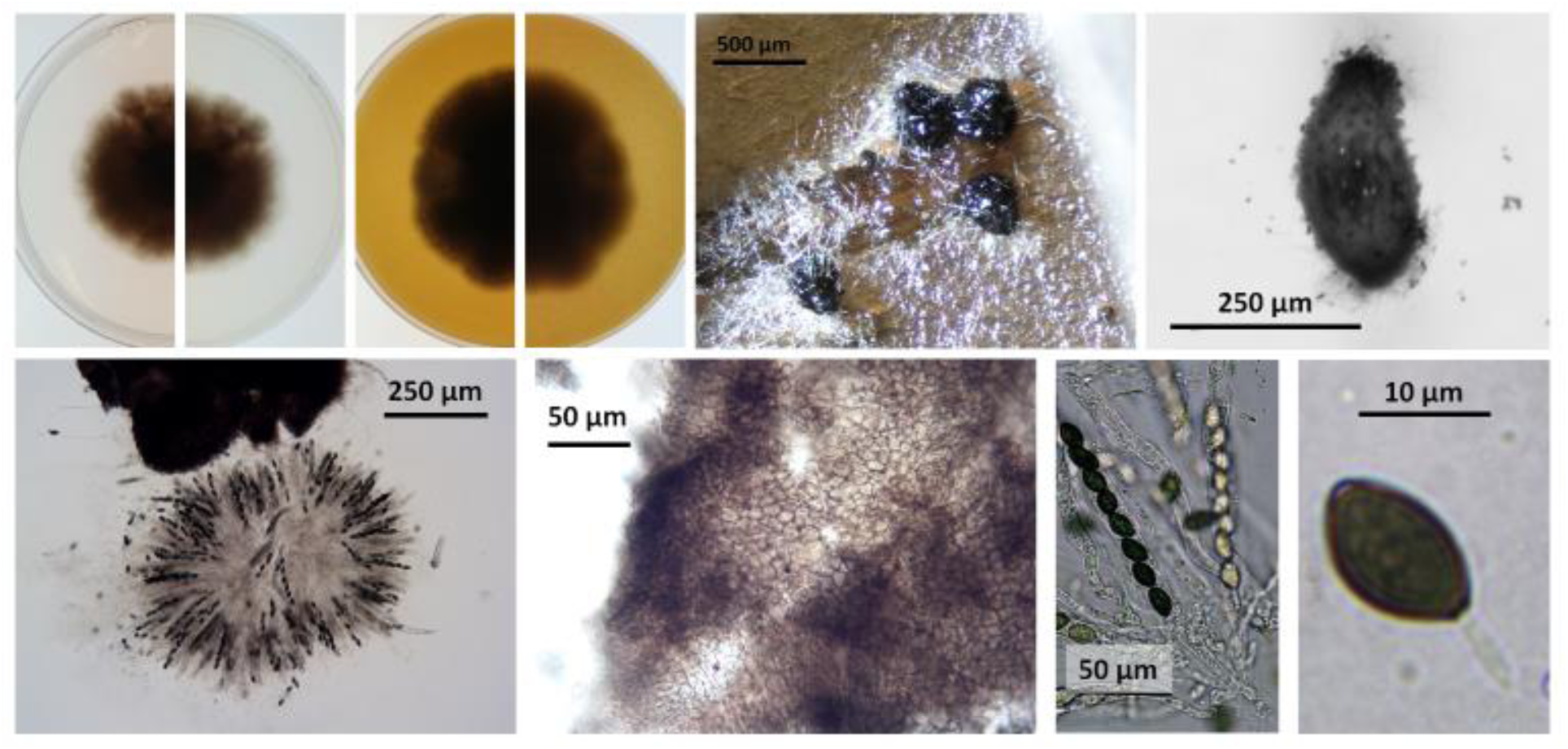
PSN1058, the ex-type of *Schizothecium octosporum*. Top from left to right: culture on M2 top view, culture on M2 reverse view, culture on V8 top view, culture on V8 reverse view, perithecia on M0+ miscanthus, isolated perithecia. Bottom from left to right; squeezed perithecium with rosette of asci and some peridium, enlarge peridium view, ascus and ascospore.

Index Fungorum number:

*Holotype*: PC0798994, ex-type: PSN1058

Description: mycelium slow growing (1.6 +/- 0.3 mm/day on M2, 2.3 +/- 0.1 mm day on V8), dark green to almost black with few aerial hyphae on M2 and profuse ones on V8 and often but not always undergoing lysis. Perithecia conical to ovoid, 600 +/- 50 x 290 +/- 40 µm, membranous, pale brown, semi-transparent when young, darker to almost black when old. Neck not-clearly differentiated, sometime slightly curved towards the light, decorated by small triangular short swollen hairs typical of *Schizothecium* perithecia. Peridium with *textura angularis*. Asci eight-spored and claviform. Spores obliquely uniseriate, slightly asymmetrical with one side flat and the other more convex, transversely septate with hyaline primary appendage and secondary appendages at both poles often difficult to see. Spore head 19.1 +/- 1.5 x 11.0 +/- 0.8 µm, ellipsoidal, smooth, flattened at the base, with an apical germ pore. Primary appendage (pedicel) cylindric, slightly tapering towards the apex, 10.1 +/- 0.4 x 1.7 +/- 0.4 µm. Secondary appendage present but evanescent and lacking on most expelled ascospores.

Habitat & Distribution: Saprobic on dung. Isolated from southern France. The only strain available for this species was isolated from rabbit dung collected in Villeneuve-lès-Maguelone, Hérault, France.

Mating strategy: heterothallic.

***Schizothecium dioctosporum*** Silar ***sp. nov.*** Fig. 23

**Fig. 23.**
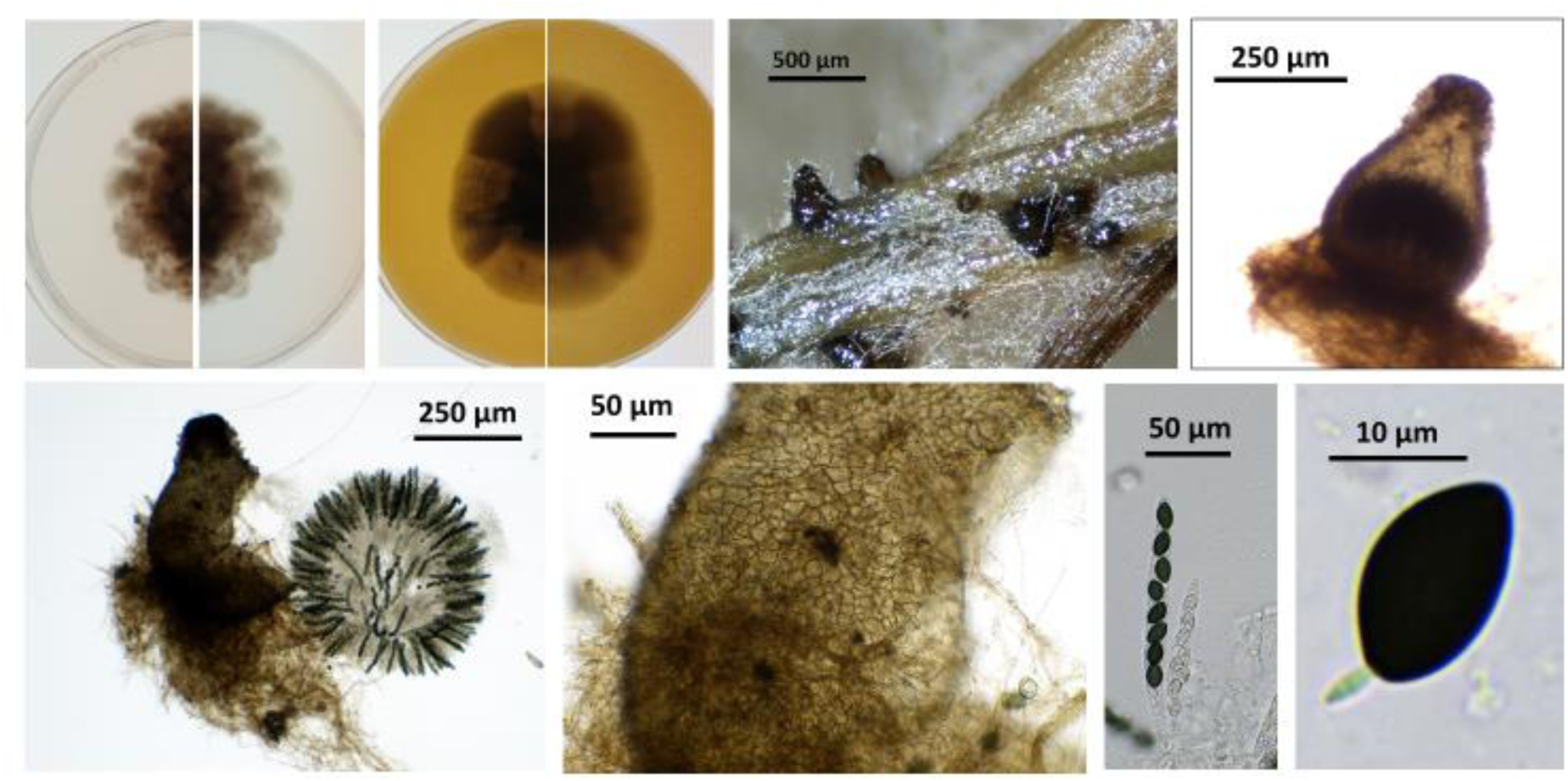
PSN1057, the ex-type of *Schizothecium dioctosporum*. Top from left to right: culture on M2 top view, culture on M2 reverse view, culture on V8 top view, culture on V8 reverse view, perithecia on M0+ miscanthus, isolated perithecia. Bottom from left to right; squeezed perithecium with rosette of asci and some peridium, enlarge peridium view, ascus and ascospore.

Index Fungorum number:

*Holotype*: PC0798993, ex-type: PSN1057

Description: mycelium slow growing (1.6 +/- 0.1 mm/day on M2, 2.3 +/- 0.0 mm day on V8), dark green to almost black with few aerial hyphae on M2 and profuse ones on V8 and often but not always undergoing lysis. Perithecia ovoid, 500 +/- 60 x 380 +/- 60 µm, membranous, pale brown, semi-transparent when young, darker when old. Neck clearly differentiated, sometime slightly curved towards the light, decorated by small triangular short swollen hairs typical of *Schizothecium* perithecia. Peridium with *textura angularis*. Asci eight-spored and claviform. Spores obliquely uniseriate, slightly asymmetrical with one side flat and the other more convex, transversely septate with hyaline primary appendage and secondary appendages at both poles often difficult to see. Spore head 19.0 +/- 0.3 x 11.0 +/- 0.1 µm, ellipsoidal, smooth, flattened at the base, with an apical germ pore. Primary appendage (pedicel) cylindric, slightly tapering towards the apex, 7.9 +/- 1.5 x 1.8 +/- 0.3 µm. Secondary appendage present but evanescent and lacking on most expelled ascospores.

Habitat & Distribution: Saprobic on dung. Isolated from southern France. The only strain available for this species was isolated from rabbit dung collected in Villeneuve-lès-Maguelone, Hérault, France.

Mating strategy: heterothallic.

## Acknowledgements

This work was supported by the EvolSexChrom ERC advanced grant #832352 (H2020 European Research Council) to T.G. and intramural funding from Université Paris Cité to P.S. The funders had no role in study design, data collection and analysis, decision to publish, or preparation of the manuscript. We thank the following collectors for samples : Pierre Defos du Rau, Yves Hurand, Mélanie Brovelli, Jacqui Shykoff, Greg Thorn, Karen Vanderwolf, Delphine Paumier, Jean Marie Ouary (association Milles Traces), Jérome Letty (Office français de la biodiversité), Paul Jay, Stella McQueen, Anne Génissel, Vénorique Decroocq, Lucas Bonometti and Pierre Gladieux. We thank Ricardo C. Rodríguez de la Vega for advices on genomic analyses and Aurélien Renault for its expert technical assistance

## Statements and Declarations

The authors have no relevant financial or non-financial interests to disclose.

All authors contributed to the study conception and design. Material preparation was mostly performed by Valérie Gautier, Emilie Levert, Elizabeth Chahine and Philippe Silar. Data collection and analysis were mostly performed by Elsa De Filippo, Christophe Lalanne, Fanny E. Hartmann, Tatiana Giraud and Philippe Silar. The first draft of the manuscript was written by Philippe Silar and all authors commented on previous versions of the manuscript. All authors read and approved the final manuscript.

The datasets generated during and/or analyzed during the current study are freely available in the GenBank repository, see Table 1 and Table S1 for accession numbers. The ex-type living cultures can be obtained in the “Centre International de Ressources Microbiennes - Champignons Filamenteux” (CIRM-CF, INRAE, France) when these were not already available in CBS or CABI collections. All the other strains are available from the corresponding author on reasonable request.

